# Multiscale computational model predicts how environmental changes and drug treatments affect microvascular remodeling in fibrotic disease

**DOI:** 10.1101/2024.03.15.585249

**Authors:** Julie Leonard-Duke, Samuel M. J. Agro, David J. Csordas, Anthony C. Bruce, Taylor G. Eggertsen, Tara N. Tavakol, Thomas H. Barker, Catherine A. Bonham, Jeffery J. Saucerman, Lakeshia J. Taite, Shayn M. Peirce

## Abstract

Investigating the molecular, cellular, and tissue-level changes caused by disease, and the effects of pharmacological treatments across these biological scales, necessitates the use of multiscale computational modeling in combination with experimentation. Many diseases dynamically alter the tissue microenvironment in ways that trigger microvascular network remodeling, which leads to the expansion or regression of microvessel networks. When microvessels undergo remodeling in idiopathic pulmonary fibrosis (IPF), functional gas exchange is impaired due to loss of alveolar structures and lung function declines. Here, we integrated a multiscale computational model with independent experiments to investigate how combinations of biomechanical and biochemical cues in IPF alter cell fate decisions leading to microvascular remodeling. Our computational model predicted that extracellular matrix (ECM) stiffening reduced microvessel area, which was accompanied by physical uncoupling of endothelial cell (ECs) and pericytes, the cells that comprise microvessels. Nintedanib, an FDA-approved drug for treating IPF, was predicted to further potentiate microvessel regression by decreasing the percentage of quiescent pericytes while increasing the percentage of pericytes undergoing pericyte-myofibroblast transition (PMT) in high ECM stiffnesses. Importantly, the model suggested that YAP/TAZ inhibition may overcome the deleterious effects of nintedanib by promoting EC-pericyte coupling and maintaining microvessel homeostasis. Overall, our combination of computational and experimental modeling can explain how cell decisions affect tissue changes during disease and in response to treatments.

## Introduction

Idiopathic pulmonary fibrosis (IPF) is a progressive and fatal fibrotic disease of the lung characterized by heterogenous formation of fibrotic foci throughout the lung. In the U.S. over 40,000 people are diagnosed with IPF annually with a mean survival rate of two to three years after diagnosis. Disease progression, quantified by lung function measurements, can be highly variable across patients, with some rapidly declining within a year of diagnosis while others decline slower across five to six years post-diagnosis^1–5^. Decline in lung function is primarily due to the thickening and stiffening of the interstitial space in the lung and loss of alveolar structures. This, in turn, has deleterious effects on the microvasculature, the network of small blood vessels, including capillaries, that are responsible for oxygen and carbon dioxide exchange^4^.

Microvascular impairments in IPF are evidenced by a decline in the diffusion capacity for carbon monoxide (DLCO) in IPF patients, which is reflective of a disruption of the alveolar-capillary membrane^6^. However, the molecular and cellular mechanisms by which microvessels are affected by IPF are not fully understood, and reports of excessive, leaky vasculature conflict with reports of vascular regression^7–10^. Both of these microvascular “phenotypes” are the result of microvascular remodeling, a process mediated by interactions between the two cells that compromise the microvasculature, endothelial cell (ECs), and pericytes, the cells that physically enwrap ECs. Proper EC-pericyte coupling maintains microvascular homeostasis^11–15^, ensuring adequate perfusion^16–18^ to enable the exchange of carbon dioxide (CO_2_) and oxygen (O_2_) across the alveolar interstitial space^19–21^ and regulating immune cell trafficking^22–28^. However, when the environment is perturbed due to disease or the introduction of a drug, cell fate decisions lead to cellular states that collectively give rise to tissue-level microvascular phenotypes. For example, in response to pro-angiogenic stimuli, ECs and pericytes temporarily uncouple from one another to allow for capillary sprouting and expansion of the microvascular network. But, ECs and pericytes subsequently recouple with one another to stabilize the newly formed capillaries^15, 29^. In contrast, when anti-angiogenic cues dominate, EC-pericyte uncoupling is typically followed by apoptosis of one or both cell types, leading to capillary regression, or the pruning of previously existing microvessels^15, 29^.

Several growth factors that mediate EC-pericyte coupling are dysregulated in IPF, including VEGF-A, PDGF-BB, and Ang 1 & 2^10, 30–36^. In patients with IPF and animal models of IPF, pericytes decouple from ECs and express myofibroblast markers, such as collagen and alpha smooth muscle actin (αSMA)^37–39^. Importantly, the two FDA approved drugs used to treat IPF, nintedanib and pirfenidone, target key signaling pathways in ECs and pericytes. Nintedanib is of particular interest in this context, as it inhibits both VEGFR, a key pro-survival and pro-angiogenic cue, and PDGFR, which is important for pericyte recruitment. Nintedanib also inhibits fibroblast growth factor receptor (FGFR), which is known to regulate angiogenesis and pericyte recruitment^40, 41^. Additionally, fibrotic lesions, or foci, in IPF undergo progressive and spatially heterogeneous mechanical stiffening of the ECM^42^, which can affect EC and pericyte dynamics^38, 43^. Understanding how dysregulated biochemical and biomechanical signals impact EC-pericyte coupling in IPF necessitates the use of a computational model where the effects of these signals can be explored, both in isolation and in combination.

Cell fate decisions are dictated by combinations of intracellular, intercellular, and environmental cues. The summation of cell fate decisions made by individual cells in a tissue results in an emergent tissue response, such as an expanded, or regressed, microvascular network. Multiscale computational models allow us to investigate how these different overlapping intracellular and extracellular cues integrate within cells to generate tissue-level outcomes, as well as to isolate how alterations in one cue affects the overall system^44^. Our group has previously developed multiscale models by combining agent-based modeling (ABM) of multiple cells in a population with logic-based ordinary differential equations modeling of intracellular signaling networks within each simulated cell^45^. We have used this framework to predict how pro-fibrotic growth factors and pro-inflammatory cytokines in the extracellular environment affect intracellular signaling within individual fibroblasts, causing them to secrete collagen and modify their extracellular environment.

Here, we present a new multiscale computational model that combines ABM^45–49^ with logic-based ordinary differential equations modeling^50–53^ to explore how biochemical and biomechanical signals integrate to affect EC and pericyte cell fate changes that give rise to structural microvascular adaptions in IPF. Intracellular signaling in individual pericytes and ECs is represented by logic-based network models implemented in Netflux^50^, and cell-cell communication, cell-environment communication, and tissue level remodeling are represented by an ABM implemented in Netlogo^54^. We used the model to predict microvascular remodeling in microenvironments with different mechanical stiffnesses that are characteristic of fibrotic foci, and experimentally validated these predictions with independent experiments. We then conducted a comprehensive set of *in silico* experiments by co-varying biomechanical and biochemical inputs, which yielded the novel prediction that nintedanib treatment in the presence of increased stiffness impairs EC-pericyte coupling by potentiating EC apoptosis and pericyte-to-myofibroblast transition (PMT). Finally, we use to the model to identify a signaling pathway that could be therapeutically targeted to mitigate the potentially harmful effects of nintedanib on EC-pericyte coupling. Our combination of multiscale computational modeling with experiments provides new insights into how cellular phenotypic states driven by biomechanical and biochemical stimuli integrate to cause tissue-level changes in disease and in response to drug treatment.

## Results

### Endothelial Cell and Pericyte Intracellular Signaling Models and Experimental Validation

Two logic-based network models of the intracellular signaling networks within ECs (**Figure 1A**) and pericytes (**Figure 1B**) that regulate cell fate decisions, such as migration, apoptosis, and quiescence, were developed in Netflux^50^. First, signaling pathways that were central to EC and pericyte cell fate decision, such as VEGFR2, a major regulator of EC quiescence and proliferation^55^, and PDGFRβ, which is a regulator of pericyte quiescence and migration^29^, were identified through literature curation and then combined with one another to construct a network diagram. Each protein, mRNA, or cell fate decision was represented as a “node” in network diagram, while the causal connections between the nodes (representing downstream signals) were represented by “edges”. The network model diagram was then converted to an ordinary differential equation-based (ODE) model using Netflux^50^. Briefly, ODE equations were solved using a normalized Hill ODE to represent the activity of each node in the model with default parameters and logic gating, thus allowing inputs to be scaled from 0 to 1. For ECs, network outputs were simplified to represent three cell fate decisions: angiogenic, apoptotic, and quiescent. These cell states collectively contribute to microvessel network (tissue-level) outcomes, including microvessel remodeling that leads to expansion of the microvascular network, microvessel remodeling that leads to reduction of the microvascular network, and homeostasis (the absence of microvessel remodeling). In total the EC model includes 32 species, 7 inputs, 32 reactions, and 3 cell fate outputs. Network outputs for the pericyte model represents three cell fate decisions: migratory, pericyte-to-myofibroblast transition (PMT), and quiescent. The pericyte model includes 32 species, 5 inputs, 34 reactions, and 3 cell fate outputs.

**Figure 1.**
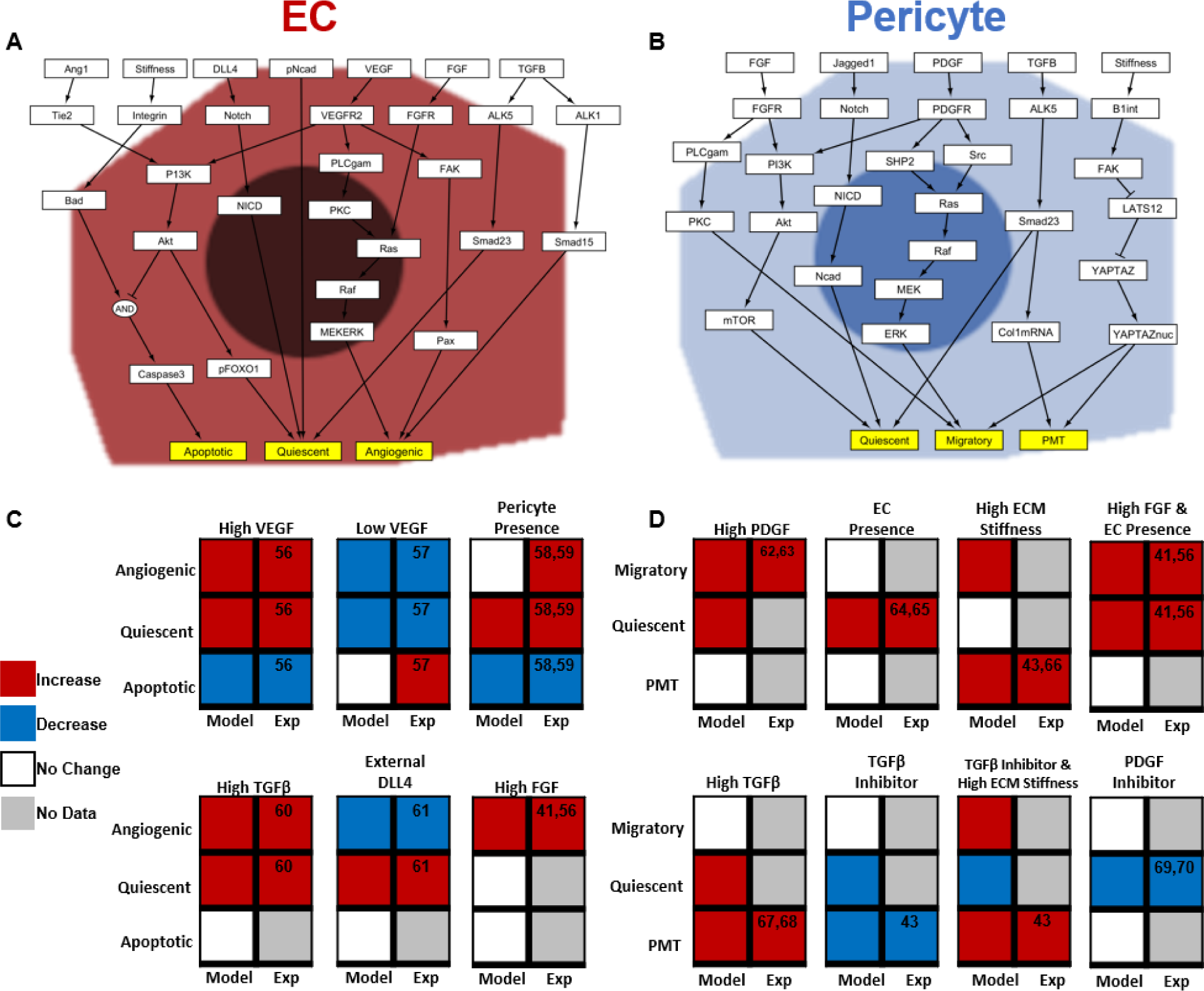
EC and pericyte intracellular network signaling model and predictions compared to published literature. (A) Schematic of the EC network model where each white box represents a specific signaling node (i.e., protein, transcription factor, or gene), and each yellow box represents a cell fate state change. (B) Schematic of the pericyte network model where each white box represents a specific signaling node (i.e., protein, transcription factor, or gene) and each yellow box represents a cell fate state change. (C) Predictions made by the EC network model (“Model” column) about the cell fate outcomes resulting from six cellular and environmental perturbations relevant to IPF, including high^56^ and low VEGF^57^, contact with a neighboring pericyte (pericyte presence)^58, 59^, high TGFβ^60^, contact with a neighboring tip EC (presence of external DLL4)^61^, and high FGF^41, 56^. Model predictions were compared to published experiments not used for model development (“Exp” column). There was agreement between model predictions and experimental outputs for 12/14, or 85%, of the tested perturbations. (D) Predictions made by the pericyte network model (“Model” column) about the cell fate outcomes resulting from eight cellular and environmental perturbations relevant to IPF, including high concentrations of PDGF-BB^62, 63^, pericyte contact with a neighboring EC^64, 65^, high ECM stiffness^43, 66^, high FGF & contact with an EC (EC presence)^41, 56^, high concentrations of TGFβ^67, 68^, TGFβ inhibitor^43^, a combination TGFβ inhibitor and high ECM stiffness^43^, and a PDGF inhibitor^69, 70^. Model predictions were compared to published experiments (“Exp” column). There was agreement between model predictions and experimental outputs for 9/9, or 100%, of the tested perturbations.

In order to validate the EC and pericyte models of intracellular signaling, predictions from both models (**Figure 1C&D**) were compared to published experiments independent from those used to develop the models. A literature search was conducted to identify publications describing an experiment with ECs, pericytes, or a combination thereof, and a perturbation of interest, such as dosing with TGFβ. The analogous simulation was then performed in Netflux by changing the input weight, *w*, of an equation from its baseline value to a value that represented the effect of the perturbation. For example, to validate the response to high levels of TGFβ in both EC and pericyte models, the input weight for TGFβ, *w*_TGFβ_, was increased from baseline (0.35) to 0.5 and changes in relative activation of the output node were quantified and compared to the baseline relative activation. These qualitative changes in node activity were then compared to their corresponding experimental findings. For example, since high TGFβ was predicted by the pericyte intracellular signaling model to increase PMT, and the corresponding experiment also demonstrated increased levels of PMT, as evidenced by αSMA and Collagen 1 expression^43^, we concluded that the model prediction matched the experiment. We further validated that under baseline conditions our models were not overly sensitive to any one input, and we explored the effect of high ECM stiffness on network sensitivity (Supplementary Figure 1).

### Mechanistic sub-network analysis suggests nintedanib regulates EC and pericyte phenotypes through similar pathways

Nintedanib is a receptor tyrosine kinase inhibitor that prevents phosphorylation of VEGFR, PDGFR, and FGFR, all three of which are expressed by ECs or pericytes and regulate microvascular homeostasis (**Figure 2A**). To explore the effects of nintedanib on ECs and pericytes individually, we performed a mechanistic sub-network analysis to identify which nodes in the model most influenced nintedanib’s effect on cell fate. This analysis was performed in two parts. First, treatment with nintedanib was simulated by reducing the activation weights, *w*, of VEGFR, PDGFR, and FGFR by 90% from 1 to 0.1 and quantifying the change in network activity. Then, a sensitivity analysis was performed by sequentially reducing each node’s weighting to zero (w_node_ = 0). The overlap of these two analyses were then determined, where nodes that both respond to nintedanib and whose knockdown affect the specified phenotype are determined to be in the “mechanistic sub-network” (Figure 2B) predominantly responsible for determining how nintedanib affects ECs and pericyte signaling and cell fate decisions in the model

For both cell types, we focused our mechanistic sub-network analysis on the cell fate decisions that are reversible, unlike apoptosis for ECs or PMT in pericytes, which are considered terminally-differentiated cell states for our model. Quiescence in both cell types was regulated by a PI3K dependent pathway, although this was activated by VEGFR in ECs and by a combination of FGFR and PDGFR in pericytes (**Figure 2C&D**). In ECs, VEGFR also regulated nintedanib’s control over the angiogenic state in conjunction with FGFR signaling, although through different pathways. FGFR regulated the angiogenic state primarily through the Ras/Raf/MEK/ERK pathway, as previously observed in the literature^71^. VEGFR, which is connected with several downstream pathways in our model leading to angiogenesis, had its strongest effect via the FAK/Paxillin pathway. For pericytes, FGFR was predicted to be the major regulator of pericyte migratory response to nintedanib.

**Figure 2.**
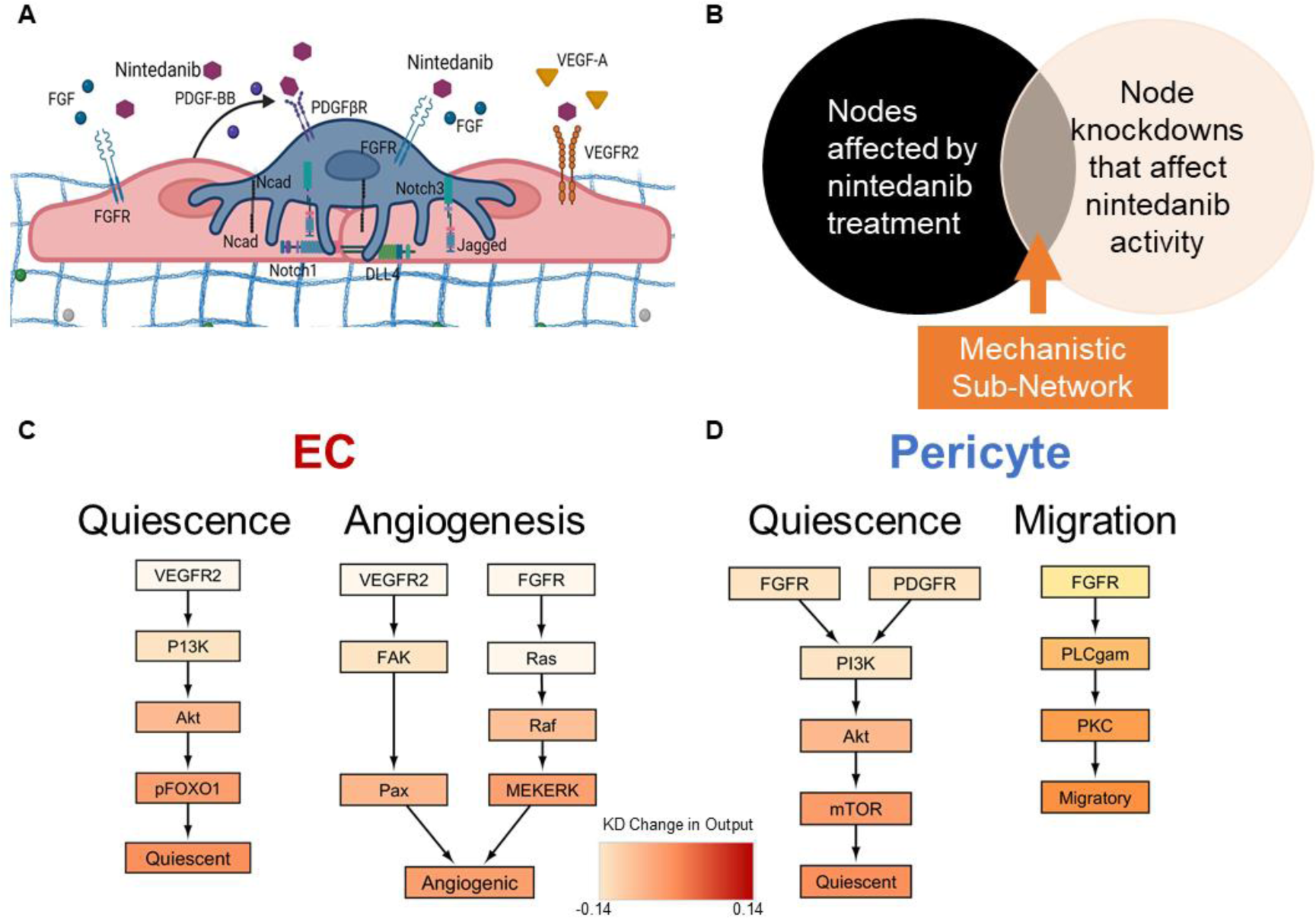
Mechanistic sub-network analysis of logic-based network models. (A) Diagram of how nintedanib targets VEGFR, PDGFR, and FGFR (B) Overview of mechanistic subnetwork analysis (C) Sub-network analysis of nintedanib regulation of EC phenotype reveals quiescence is regulated in a VEGFR dependent manner alone while angiogenesis is controlled by both VEGFR and FGFR. (D) Sub-network analysis of pericyte response to nintedanib reveals FGFR is key regulator of both quiescence and migration in pericytes while PDGFR plays a strong role in maintaining pericyte quiescence. Sub-figure A was created using Biorender.com.

### Validation of the multiscale model’s ability to predict microvascular remodeling in response to altered levels of mechanical stiffness in the microenvironment

A multiscale computational model of the lung microvasculature was constructed by connecting both the EC and pericyte intracellular signaling network Netflux models described above to a 2-dimensional (2D) ABM of a lung microenvironment, where the locations of simulated ECs and pericytes were approximated from an immunofluorescence (IF) image of healthy lung microvasculature (**Figure 3A**). The ABM, representing 500μm x 500μm slice of lung tissue in a 41 x 41 square grid of pixels (with each pixel representing 12 x 12 microns), was built in NetLogo^54^. The ABM contains information about the extracellular cues, such as ECM stiffness and growth factor concentrations, at each pixel location in the 2D model (**Figure 3B**). The Py extension in NetLogo was used to connect the ABM model with the two Netflux models thereby generating a multiscale model that integrates intracellular cues and cell fate decisions (**Figure 3C**). These intracellular cues and cell fate decisions were computed by the Netflux models for each simulated cell while tissue properties were computed by the ABM to predict changes in microvascular network architectures and population-level cell fate outcomes, such as number of apoptotic ECs or percent of pericytes that have undergone PMT.

**Figure 3.**
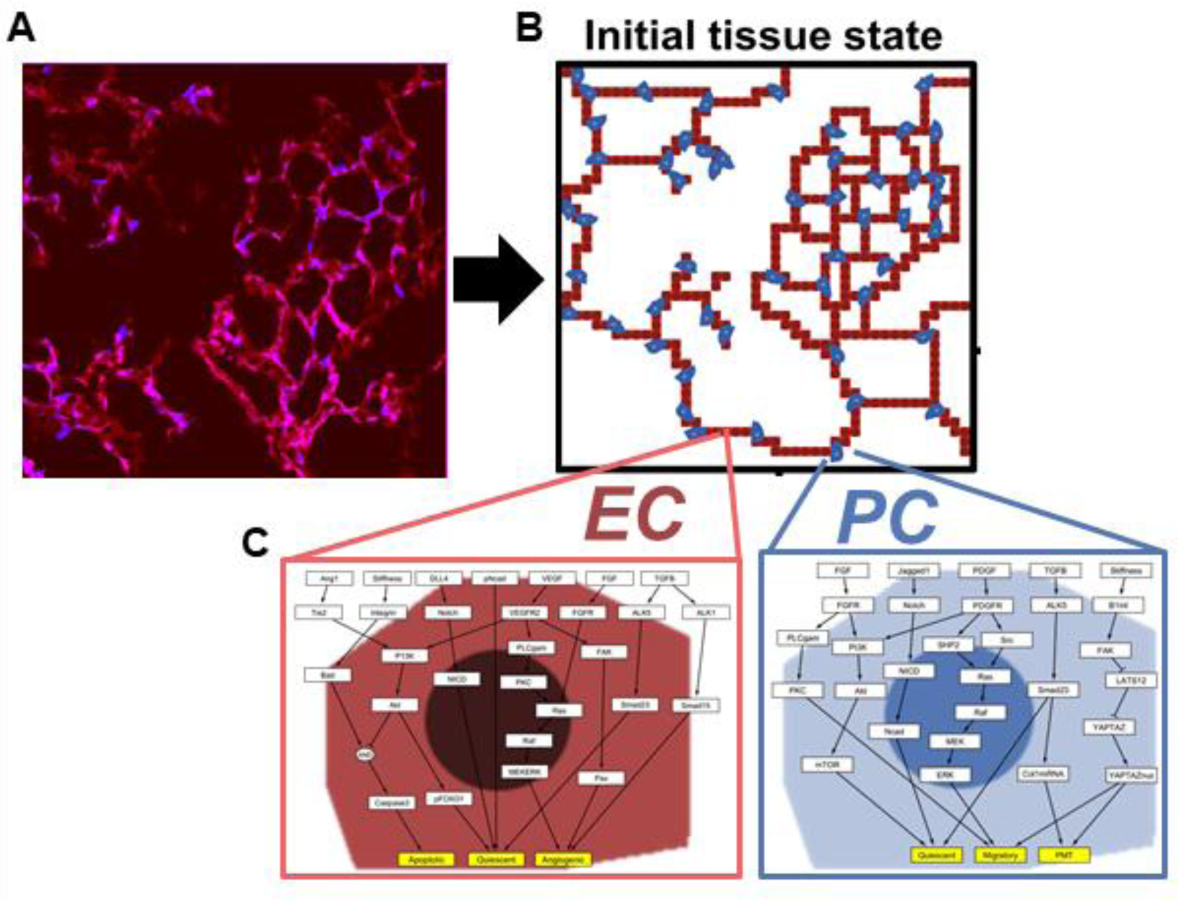
Overview of multiscale model framework. (A) An IF image of healthy lung was use to prescribe initial locations for simulated ECs (red, CD31) and pericytes (purple, TdTomato Myh11 Cre labelled) in the ABM environment in NetLogo (B). (C) The logic-based network models representing ECs and pericytes were connected to the ABM environment using the Py extension in NetLogo.

At each time step in the ABM representing 6 hours, the cellular agents sense and respond to their local extracellular cues, and signals from neighboring cells, and input them into the logic-based network model. For each cell agent, the network model computes the relative activation of the cell fate decision nodes, and the cell fate decision that has the highest relative activation is enacted by that cell in the ABM. Simulated ECs are able to be in an angiogenic, apoptotic, or quiescent state, and pericytes are able to remain quiescent, migrate, or undergo pericyte-to-myofibroblast transition. Over the entire simulation time course (representing seven days), microvessel network (tissue-level) outcomes emerge as a summation of these independent cell decisions.

To validate the multiscale model’s ability to predict microvessel network remodeling in response to microenvironmental stiffnesses variations, we simulated the responses of ECs and pericytes to stiffnesses ranging from 2 kPa to 20 kPa recapitulating the stiffness gradient seen from healthy to IPF lung^42^. The multiscale model predicted that as ECM stiffness increases from healthy lung (2 kPa), to immature fibrosis (10 kPa), and mature fibrosis (20 kPa), the extent of the microvessel network, quantified by vessel area, significantly decreases (**Figure 4A**), and this was associated with significant increases in EC apoptosis (**Figure 4B**). The mean vessel area at 2 kPa, 48.2%±0.7, was higher than the initial vessel area of 23.5%, and the mean number of ECs that underwent apoptosis was 8.7±2.8, indicating that some levels of homeostatic remodeling occurred (**Figure 4C**). At 10kPa, mean vessel area was 44.8%±2.1, and mean number of apoptotic ECs increased to 149±18.7. This indicates that some level of angiogenesis occurred to offset the microvessel regression caused by EC apoptosis. Contrastingly, at 20 kPa (mature fibrosis), the mean vessel area dropped to 9.8%±0.58, and the average number of apoptotic ECs increased to 228.3±9.8, resulting in almost complete regression of the vascular network (**Figure 4D**).

**Figure 4.**
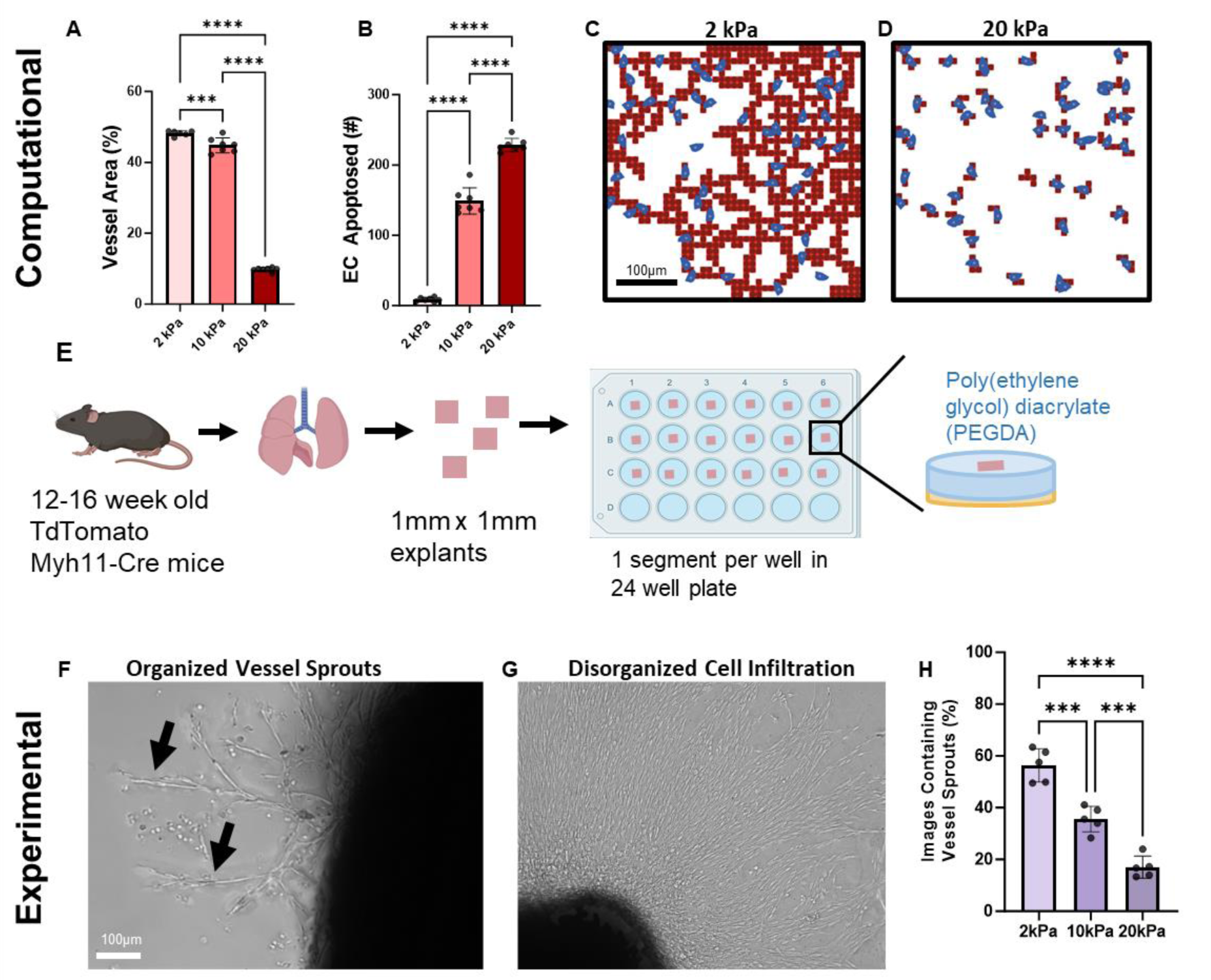
Validation of multiscale model response to increasing stiffness. Computational Results (A-D) and Experimental Results (E-H). (A) At Day 7, Percent vessel area in the multiscale computational model output decreases as stiffness increases. (B) The number of apoptotic endothelial cells in the multiscale computational model increases as stiffness increases. (C) Representative output of the multiscale computational model at 2kPa. (D) Representative output of the multiscale computational model at 20kPa. (E) Overview of experimental setup for lung explant assay (F) Representative image from the lung explant model demonstrating organized microvessel sprouts (black arrows). (G) Representative image from the lung explant model showing disorganized cell infiltration. (H) Quantification of the number of images containing organized microvessel sprouts after 7 days. Statistics: One-Way ANOVA with Tukey’s post-hoc test, ***p<0.001, ****p<0.0001. Parts of (E) were created using Biorender.com.

To determine if these predictions were biologically accurate, we utilized a murine lung explant angiogenesis assay that we have previously developed^72^. Lungs from mice expressing an endogenous pericyte lineage-tracing fluorescent reporter (abbreviated, TdTomato Myh11-Cre mice, and described in the Methods) were sliced into 1 mm x 1mm x 1 mm explants and then individually plated onto 6 mm diameter PEGDA hydrogels containing a combination of PDGF-BB and FGF-2 and with one of three stiffnesses matching those simulated in the computational model: 2 kPa, 10 kPa, and 20 kPa (**Figure 4E**). After 7 days of culture, the explants were imaged and scored based on the presence of organized microvessel sprouts (**Figure 4F**) or disorganized cell infiltration (**Figure 4G**), as we have published previously^72^. As hydrogel stiffness increased, the percent of images containing microvessel sprouts significantly decreased. This result correlated with our multiscale computational modeling results indicating that increased ECM leads to a decrease in vascular structures (**Figure 4H**).

### Predicting Changes in Cell Fate Decisions Over Time in Response to ECM Stiffness

One advantage of having a multiscale model is that we can explore how the underlying cell fate decisions that drive microvascular remodeling in different environments change over time. We quantified how many ECs and pericytes were predicted in each cell fate category for each step of our model for 2 kPa, 10 kPa, and 20 kPa. At 2 kPa, we observed an initial drop in the percentage of quiescent ECs that was accompanied by a spike in the number of angiogenic cells. However, the overall system returned to a steady state of 95% quiescent cells at approximately Day 3 (**Figure 5A**). Over 95% of pericytes in these simulations maintained a quiescent state with small spikes in the percent of migratory pericytes in the earlier time points, aligning with the return of higher levels of quiescence in ECs (**Figure 5B**). In 10 kPa, we saw a similar disruption in the quiescent EC population but in this scenario, we observed both angiogenic ECs and apoptotic ECs, indicating variable responses to local environmental cues (**Figure 5C**). The return of 95% of the ECs to a quiescent cell state occurred on Day 6. In the 10 kPa microenvironment, we observed higher levels of migratory pericytes throughout the simulation, but over 95% of all pericytes maintained a quiescent steady state (**Figure 5D**). In the 20 kPa microenvironment, the reduction in the percentage of quiescent ECs was accompanied by a large spike in the percentage of apoptotic ECs. However, the system quickly returned to a 95% quiescent cell steady state after one day, with occasional small spikes in apoptotic EC over the time course (**Figure 5E**). Conversely, by Day 3 fewer than 95% of pericytes were in a quiescent state, and they continued to decline over the simulation time course (**Figure 5F**). This decline was accompanied by an increase in the percentage of migratory pericytes. Importantly, we also began to see low but persistent percentages of pericytes undergoing PMT at Day 4.5.

**Figure 5.**
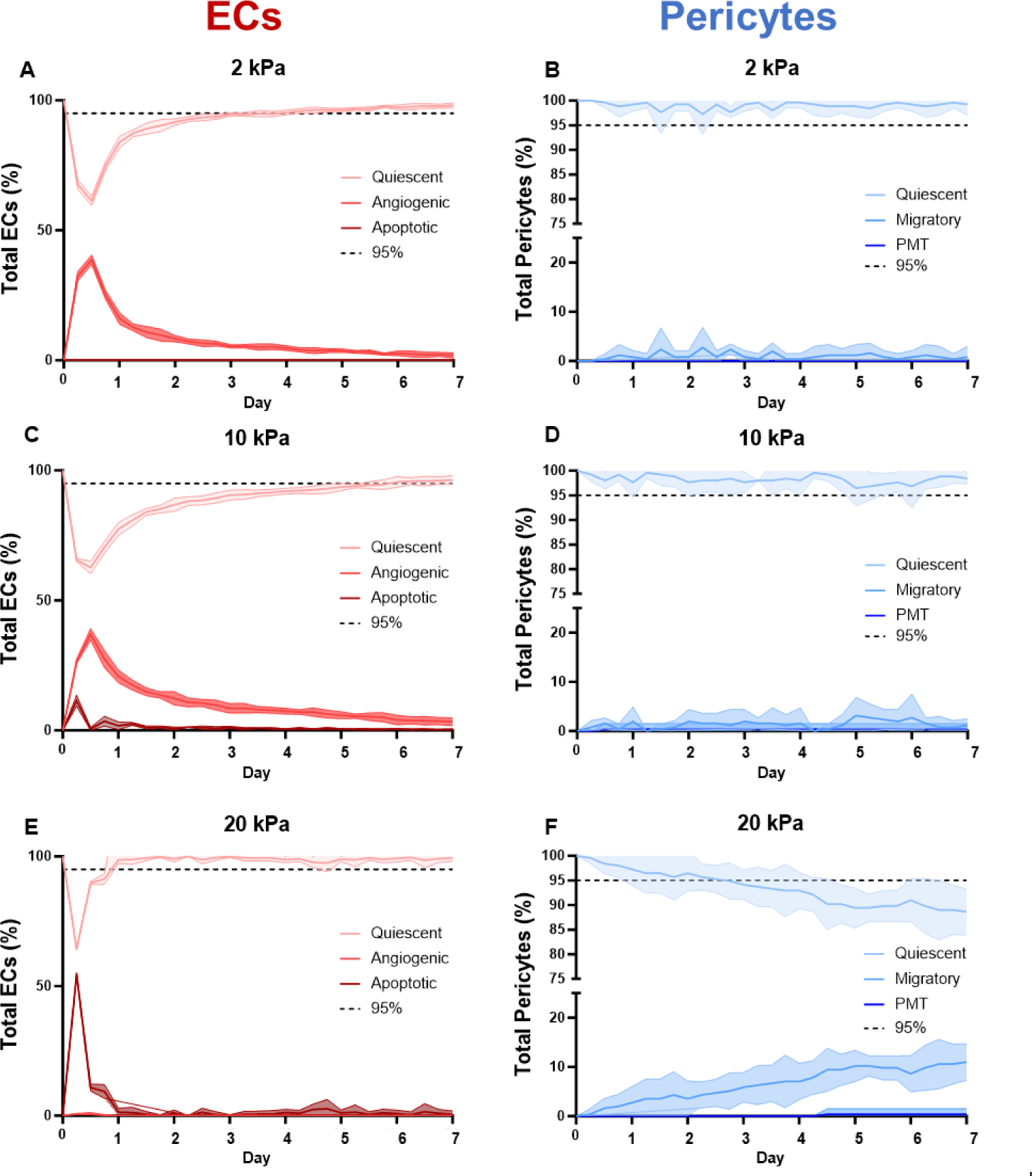
The effect of ECM stiffness on cell fate decisions over time. (A) At 2 kPa an initial decrease in the percentage of quiescent ECs is accompanied by an increase in angiogenic ECs before returning to a 95% quiescent steady state. (B) The pericytes at 2 kPa overall maintain a steady state of over 95% quiescent, with the occasional spike in the percent of migratory pericytes. (C) At 10 kPa the initial drop in quiescent ECs was accompanied by an increase in both angiogenic and apoptotic ECs resulting in a delayed response to 95% quiescent steady state. (D) At 10 kPa there is still an overall pericyte steady state of above 95% quiescent cells but with an increase in the occurrence of migratory pericytes. (E) At 20 kPa a sharp decline in quiescent ECs is due to an increase in apoptosis but a quick return to a 95% quiescent steady state. (F) The percent of quiescent pericytes declines throughout the time course at 20 kPa and is associated with an increase in the percent of migratory pericytes and the occurrence of PMT. N = 5, shading represents 95% confidence interval.

### Predicting microvascular remodeling in variable stiffness environments in response to nintedanib

We next used the multiscale model to evaluate how treatment with nintedanib, 1 of 2 drugs FDA-approved for the treatment of IPF, affected microvascular remodeling in microenvironments with mechanical stiffnesses of 2 kPa, 10kPa, and 20 kPa. Nintedanib inhibits VEGFR on ECs, PDGFR on pericytes, and FGFR, which is expressed by both cell types. Treatment with nintedanib was modeled by reducing the maximum activation weight of each receptor to 0.1 in both the EC and pericyte models. Based on prior literature^73^ we hypothesized that nintedanib would decrease vessel area in the computational model and microvessel infiltration into the hydrogel of the experimental model. Treatment with nintedanib resulted in significantly decreased vessel area in two of the three treated groups when compared to the untreated controls (**Figure 6A**). At 2 kPa, significant decreases in vessel area (48.2%±0.7 vs 44.1%±2.1) were measured, although this was not associated with an increase in the number of apoptotic ECs (8.7±2.8 vs. 6.7±3.7) (**Figure 6B**). At 10 kPa, treatment with nintedanib reduced mean vessel area to 41.6%±2.1 compared to the control (44.8%±2.1). Limited efficacy of nintedanib was observed at 20 kPa, as both groups had vessel areas of about 10% and no significant differences in number of apoptotic ECs.

**Figure 6.**
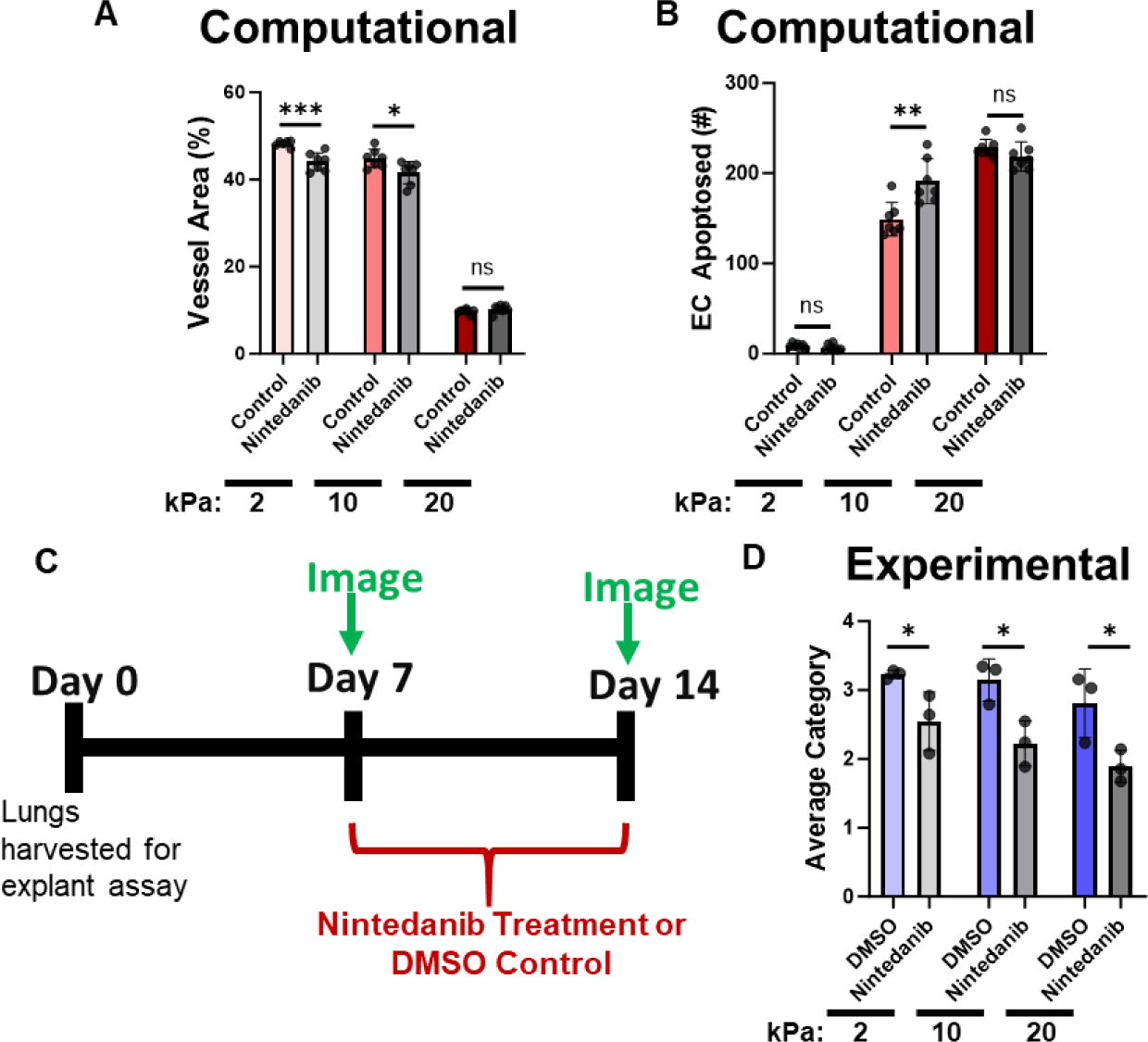
Multiscale model response to nintedanib. (A) Nintedanib significantly decreased vessel area in the multiscale model at 2 kPa and 10 kPa but had no effect at 20 kPa. (B) Number of ECs that apoptosed was significantly increased at 10 kPa when treated with nintedanib, but not in 2 or 20 kPa. (C) Experimental model timeline, after the first experimental test data on stiffness alone was gathered on Day 7, explants were treated with either 50 nM nintedanib or DMSO for seven days then imaged again. (D) Images taken on Day 14 were categorized based on vessel infiltration into gel. Nintedanib significantly decreased the average category score of vessel infiltration in all three conditions indicating less vessel infiltration into the gel. Computational Model N = 7. Experimental Model N = 3. Statistics: One-tailed t-test, *p<0.05, **p<0.01, ***p<0.001, ns = not significant.

Using the murine lung explant experimental model described above, we performed new experiments to determine if these model predictions were biologically accurate. After allowing sprouts to form for seven days, lung explants were either treated with 50 nM nintedanib in 0.05% DMSO or 0.05% DMSO control (**Figure 6C**). Seven days later, explants were imaged again and scored blindly on a scale of 1 to 5 based on the amount of vessel infiltration into the surrounding gel. The absence of cell and/or vessel structures was classified as “Category 1”, and the presence of extensive vessel structures and cell infiltration was classified as “Category 5”. At 2 kPa, the average infiltration category score across all explants treated with nintedanib was 2.55±0.42 which was significantly lower than the DMSO-treated control group (3.16±0.062) (**Figure 6D**). Similar results were observed at 10 kPa and 20 kPa, with the average infiltration category of the nintedanib-treated group (2.22±0.33 in 10 kPa and 1.89±0.23 in 20 kPa) being significantly lower than the DMSO treated control (3.144±0.31 in 10 kPa and 2.81±0.5 in 20 kPa).

### Multiscale model predicts that nintedanib disrupts EC-Pericyte coupling

EC and pericyte fate decisions at every time step were recorded for the simulations of nintedanib treatment in the microenvironments with different stiffnesses. While the EC cell fate dynamics were similar to those in the drug-free simulations, the pericyte cell fate dynamics in the presence of nintedanib demonstrated a disruption in their coupling with ECs that was not predicted in the absence of drug.

In the 2 kPa microenvironment, the multiscale model predicted the same initial, transient drop below 95% in the percentage of quiescent ECs. The transient increase in angiogenesis that the model predicted without drug, also dissipated by Day 3 in the presence of nintedanib (**Figure 7A**). The pericyte dynamics in 2 kPa with nintedanib treatment were also similar to those in the absence of drug, with 95% of pericytes remaining in a quiescent steady state for the entire simulation duration (**Figure 7B**). However, in microenvironmental stiffnesses of 10 kPa, we observed the first signs of impaired EC-pericyte coupling. While 95% of nintedanib-treated ECs returned to a quiescent state by Day 5, slightly earlier than the drug-free simulations, the percentage of quiescent pericytes fell below 95% on Day 4 and continued to decline over time (**Figure 7C&D**). This loss of quiescence in pericytes was accompanied by an increase in PMT that was not seen in the absence of drug. Finally, in the 20 kPa microenvironment, the EC dynamics were predicted to be similar in both drug and drug-free simulations, with a spike in apoptosis and rapid loss of EC quiescence that were largely reversed by Day 1(**Figure 7E**). Contrastingly, the model predicted a rapid decline in the percentage of quiescent pericytes, which dropped below 95% as early as Day 1 (compared to Day 3 in the drug-free simulations) (**Figure 7F**). This decline was concomitant with an increase in the percentage of migratory pericytes and earlier instances of pericytes adopting a PMT fate (at Day 1 as opposed to Day 4.5 in the drug-free simulation). Because nintedanib targets three pathways we wanted to dissect the role of the inhibition of each receptor in nintedanib-mediated loss of EC-pericyte coupling and vessel area (**Supplementary Figure 2**). In summary, treatment with a PDGFR inhibitor alone may be an ideal therapeutic strategy as it preserves EC-pericyte coupling at 10 kPa, delays the occurrence of PMT at 20 kPa, and prevents a stiffness-induced decrease in vessel area at 10 kPa.

**Figure 7.**
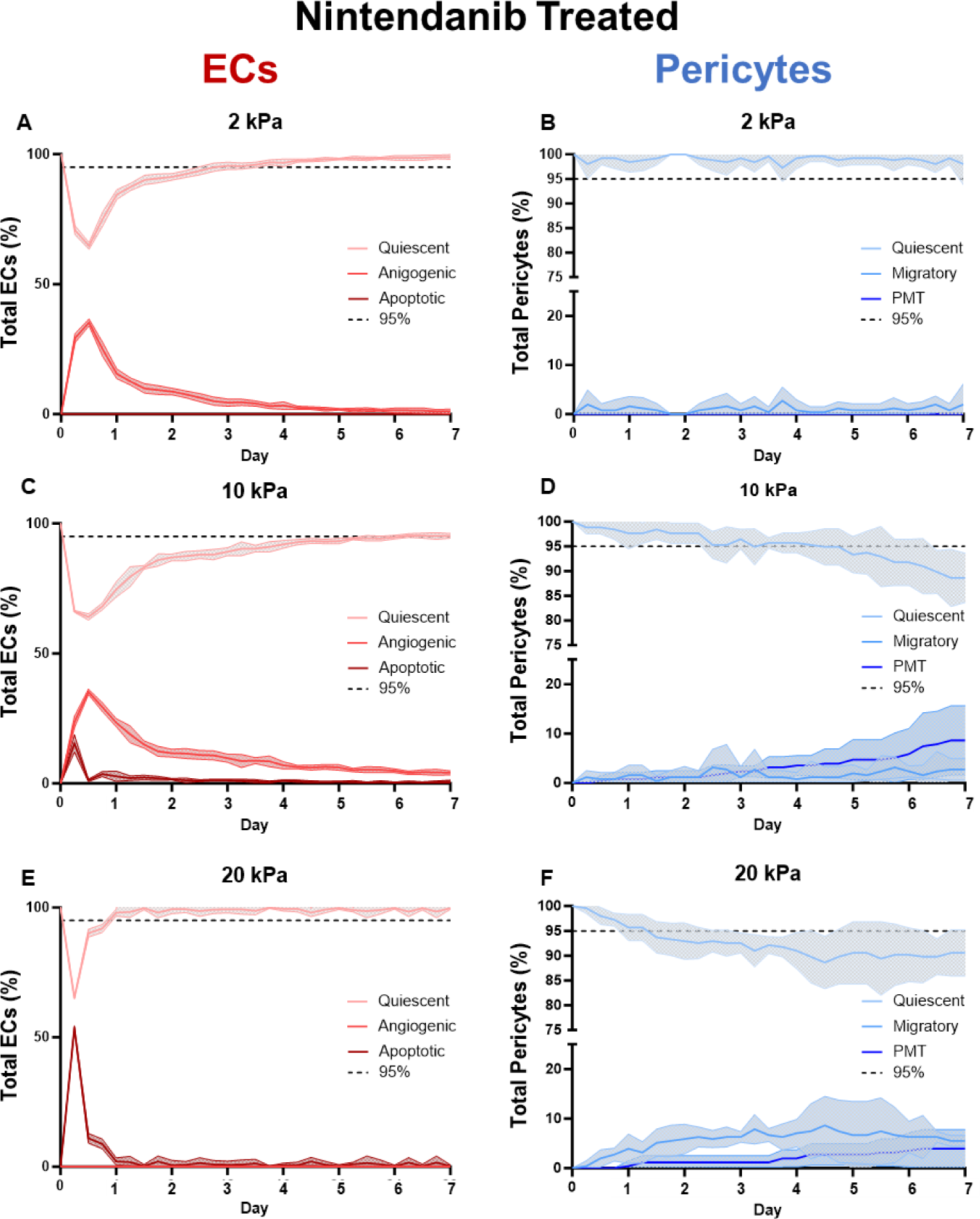
Predicted cellular states following nintedanib treatment. (A) At 2 kPa, there is an initial reduction (below 95%) in the percentage of ECs that are in a quiescent state, but by Day 3 the percentage of ECs in a quiescent state returned to 95% or greater. (B) Pericytes at 2 kPa maintain a similar level of quiescence, with the mean remaining above the 95% quiescent threshold. (C) At 10 kPa the initial drop in the percentage of quiescent ECs was accompanied by an increase in both angiogenic and apoptotic ECs, but returned to steady state on Day 5. (D) At 10 kPa there is a decline in pericyte quiescent overtime combined with an increase in migratory pericytes and pericytes undergoing PMT. (E) At 20 kPa there is a sharp decline in quiescent ECs and increase in apoptotic ECs but the cells quickly return to a 95% quiescent steady state at Day 1. (F) The percentage of quiescent pericytes declines rapidly, dropping below 95%, starting on Day 1. On Day 1 pericytes are predicted to begin exhibiting a PMT state. N = 5, shading represents 95% confidence interval.

### Multiscale model predicts a strategy to maintain EC-pericyte coupling in the presence of nintedanib

To identify a signaling pathway that could be targeted to mitigate the loss of vessel area caused by a stiff ECM and the increased percentage of pericytes exhibiting migratory and PMT states invoked by nintedanib treatment, we used the model to simulate inhibition of YAP/TAZ in the presence of nintedanib. In our model, the YAP/TAZ pathway is only present in pericytes and is upstream of both the migration and PMT cell state nodes. The pericyte network model was also found to be sensitive to YAP/TAZ knock out in the sensitivity analysis in Supplemental Figure 1. In the simulated 10 kPa microenvironment, treatment with both nintedanib and a YAP/TAZ inhibitor had no obvious effect on EC cell fate decisions (**Figure 8A**); but the percentage of quiescent pericytes remained above 95% for the full seven-day time course (**Figure 8B**). This is in contrast to the pericyte response to nintedanib alone (**Figure 7D**), wherein the percentage of quiescent pericytes gradually decreased, starting on Day 1, and dropped below 95% on Day 5. In the simulated 20 kPa microenvironment, inhibition of YAP/TAZ decreased the height of the initial spike in the percentage of apoptotic ECs at Day 0.5 (**Figure 8C**) by approximately 20% compared to treatment with nintedanib alone (**Figure 7E**). This was accompanied by a 1.5-day delay in the transition of pericyte states from the quiescent state to PMT and migratory states, which occurred at Day 2.5 with the addition of YAP/TAZ inhibition (**Figure 8D**) compared to Day 1 in the nintedanib-alone simulation (**Figure 7F**). Another difference caused by inhibition of YAP/TAZ was that the reduction in the percentage of pericytes in a quiescent state in the first half of the time course was associated with an increase in pericytes undergoing PMT. Contrastingly, this decrease in the nintedanib-alone simulation was more attributed to an increase in pericytes adopting a migratory state. The multiscale model predicted beneficial effects of these cell state changes on microvessel remodeling (**Figure 8E**). By increasing the percentage of pericytes in a quiescent state in the simulated 10 kPa microenvironment, YAP/TAZ inhibition maintained microvessel area at levels that were similar to those in the 2 kPa microenvironment. However, this preservation of microvessel homeostasis with YAP/TAZ inhibition was not observed in the 20 kPa microenvironment. We also evaluated the effect of YAP/TAZ inhibition independent of nintedanib treatment and found it was able to maintain pericyte quiescence levels above 95% at all three simulated stiffnesses (Supplementary Figure 3).

**Figure 8.**
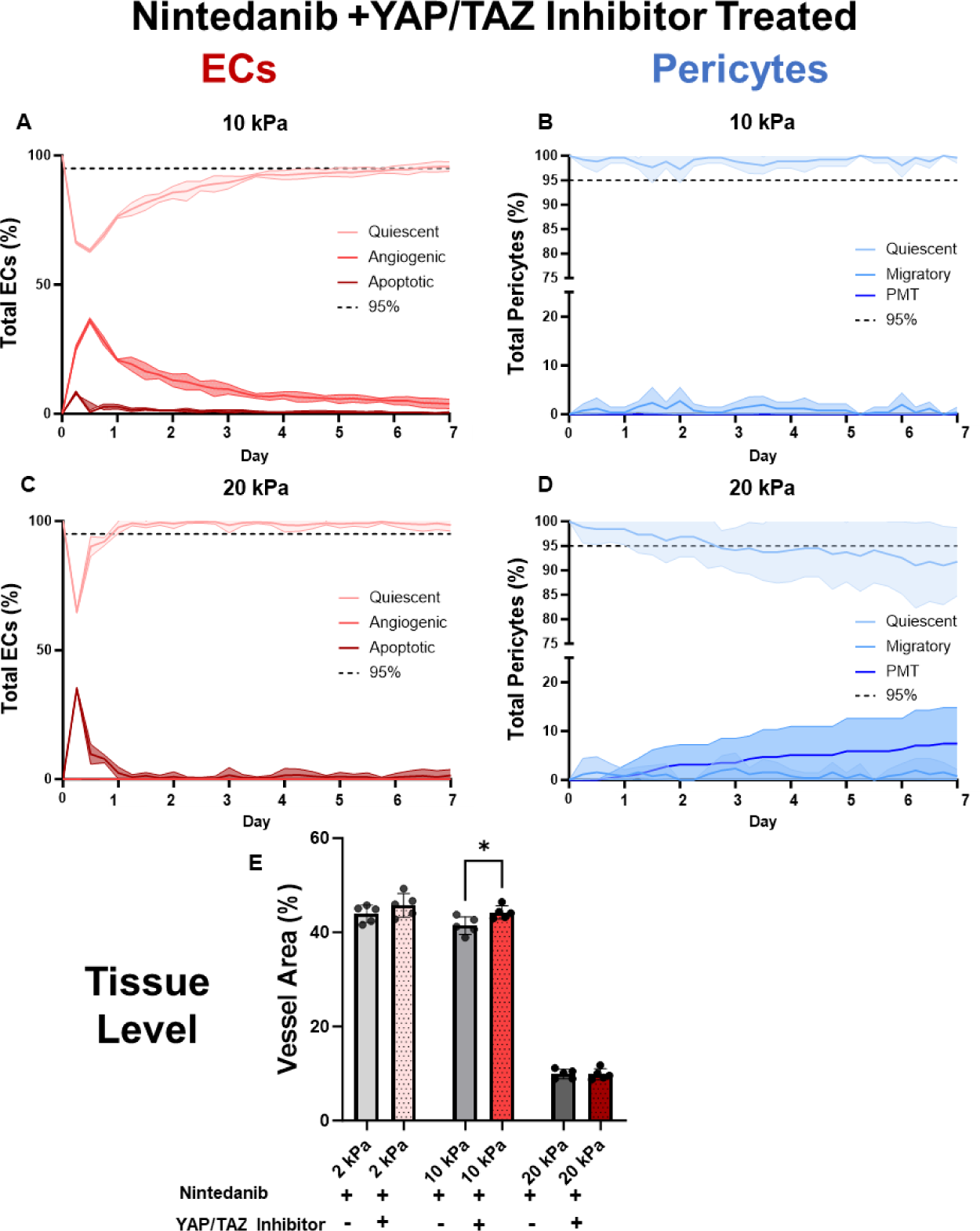
Effectiveness of combination treatment of nintedanib + YAP/TAZ inhibitor on cell- and tissue-level outcomes. (A) At 10 kPa treatment with a combination of nintedanib and a YAP/TAZ inhibitor results in a initial drop in EC quiescence that is associated with a spike in the percent of angiogenic and apoptotic ECs before eventually returning to 95% quiescent steady state at Day 6. (B) Combination treatment with nintedanib and YAP/TAZ prevents nintedanib associated loss of 95% pericyte quiescent steady state and PMT at 10 kPa. (C) At 20 kPa, combination treatment with nintedanib and a YAP/TAZ inhibitor causes an initial drop in EC quiescence that is recovered to a 95% quiescent steady state at Day 1. (D) Combination treatment delays the loss of 95% pericyte quiescence to Day 2.5 compared to the nintedanib treated alone. However, this is associated with a large influx of PMT. (E) Quantification of resulting tissue level phenotype demonstrates that by preserving pericyte quiescence in 10 kPa, the loss of vessel area due to nintedanib treatment has been recovered. Statistics: N = 5, shading represents 95% confidence interval, bar graph: two-tailed t-test.

## Discussion

Understanding mechanisms of disease and drugs across spatial scales – from molecule to tissue – remains a major challenge in biomedical research that multiscale mechanistic modeling can help overcome. We demonstrate that a multiscale model can predict how the disease microenvironment and pharmaceuticals will interact, resulting in cell fate changes that affect tissue-level remodeling. We focused our study IPF, where the impact of ECM stiffness on EC-pericyte coupling and microvascular remodeling are ambiguous, with conflicting reports of excess, leaky vasculature and regions void of any vasculature^10, 74–76^. Our multiscale model’s predictions of microvascular remodeling in response to altered mechanical stiffness alone, and in combination with nintedanib treatment aligned with our experiments. Simulations suggested a new effect of nintedanib, which prevents EC-pericyte coupling in stiffer extracellular environments (10 and 20 kPa), thereby augmenting microvessel regression. These results align with previous studies that observed decreased vessel area after nintedanib treatment in the bleomycin mouse model of IPF^77^ and reduced association of ECs and mural cells in an co-culture model treated with nintedanib^73^. Furthermore, when we used the multiscale model to identify a signaling pathway that can be targeted to preserve EC-pericyte coupling in stiffer environments, it suggested that inhibition of YAP/TAZ signaling can counteract the negative effects of nintedanib on EC-pericyte coupling prior to mature scar formation.

We started with a focus on the disease environment by using the multiscale model to investigate how the summation of individual EC and pericyte fate decisions in different microenvironmental stiffnesses impacted microvessel remodeling. At the cellular level, the model predicted how increasing ECM stiffness from 2 to 20 kPa contributed to a breakdown in communication between ECs and pericytes, as more ECs become apoptotic and more pericytes exhibited a migratory cell state. The model also predicted an increase in the percentage of pericytes undergoing PMT in the 20 kPa environment. This corresponded with our explant experimental results wherein stiff PEGDA hydrogels were sufficient to disrupt vessel sprouts. While stiffness is just one aspect of the dysregulated microenvironment in IPF, according to recent studies, a stiff ECM alone can cause vascular regression. For example, a recent study co-cultured ECs and fibroblasts in a vessel-on-a-chip model of microvasculature and showed that a stiffness of 20 kPa reduced vessel area and promoted PMT, as quantified by the expression of collagen 1 and αSMA^43^. Further, studies analyzing IPF samples report that mature scar is completely void of microvessels^74–76^, which is consistent with the extreme reduction in microvessel area in mature fibrosis (20 kPa) that was predicted by our model and validated by our experiments. Hence, our model predictions and experiments contribute to the published literature by suggesting that ECM stiffness drives EC-pericyte uncoupling, leading to EC apoptosis and pericyte PMT and causing microvascular regression.

Since nintedanib is a standard of care treatment for IPF^4^, it is important to understand how the cell state changes evoked by this drug scale-up to impact microvessel remodeling. At the intracellular level, our mechanistic sub-network analysis identified several pathways that were pertinent to how nintedanib functions inside ECs and pericytes. Of particular interest was the result that the PI3K pathway regulated quiescence in both ECs and pericytes, as a recent study in the bleomycin mouse model found that nintedanib ameliorated inflammation and fibrosis specifically through the PI3K pathway^78^. This supports the role of nintedanib in controlling cell fate through the PI3K pathway; however, our model predicted that it negatively impacts EC-pericyte coupling. Specifically, the multiscale model predicted that nintedanib disrupts the ability for EC-pericyte recoupling after an initial decrease in the percent of quiescent pericytes, potentiating PMT in environments with higher ECM stiffnesses. This prediction is important because decreases in the percentage of quiescent pericytes, even when 95% of ECs (or more) are able to return to a quiescent state, could cause excessive vascular leakage, an outcome of defective microvessel remodeling that has been reported in IPF patients^7, 8^. Excessive vascular leakage, particularly that which causes alveolar edema, can also be detrimental to gas exchange and lead to poor outcomes^79, 80^.

At the tissue level, nintedanib reduced vessel area in both the soft (2 kPa) and immature fibrotic (10 kPa) environments of our experimental model, which was consistent with our multiscale model’s predictions of vessel regression in response to nintedanib in these environments. This adds to existing literature that has demonstrated nintedanib’s anti-proliferative effects in ECs and pericytes^81, 82^, as well as inhibition of EC-mural cell coupling^73^. Importantly IPF is a progressive disease in which the lung becomes increasingly stiff over time, and characterized by significant spatial heterogeneity in fibrotic foci distribution and gene expression^83^. This suggests that the effects of currently available antifibrotic treatments vary, both temporally within an individual over time and spatially within the lung. In conclusion, nintedanib, which inhibits key pathways of communication between ECs and pericytes, exacerbates ECM stiffness-induced increases in PMT, thus impairing EC-pericyte recoupling and augmenting microvascular regression.

Finally, we used our model to determine if targeting a specific node in the intracellular signaling network of pericytes would prevent nintedanib-induced PMT, restore EC-pericyte coupling, and increase microvessel homeostasis. We chose to inhibit YAP/TAZ, which was identified as a regulator of migration and PMT in pericytes in our sensitivity analysis and has previously been studied as a target for preventing fibrosis^84, 85^. Simulating dual treatment with nintedanib and YAP/TAZ inhibition at 10 kPa improved both cell and tissue-level outcomes. The percentage of pericytes in a quiescent state remained above 95% for the entire seven-day period, maintaining EC-pericyte coupling, and promoting microvessel homeostasis. The benefits of inhibiting YAP/TAZ signaling were partially negated by an ECM stiffness of 20 kPa. Despite this, the reduction in the percentage of quiescent pericytes below 95% was delayed long enough to mitigate the initial EC apoptosis observed in this mature scar environment. Interestingly, YAP/TAZ inhibitors have been suggested to have potential benefits in IPF by mitigating fibroblast activation^84, 85^. Here, we suggest the strategy of inhibiting YAP/TAZ in combination with nintedanib to promote microvascular homeostasis.

Overall, multiscale computational modeling offers a unique framework for integrating data sets across cell, tissue, and organ-level contexts. In the present study we combine logic-based network modeling, which captures complex intracellular signaling networks and pharmacological responses at the single-cell level, with an ABM that provides a multi-cell, spatial and stochastic framework^86^. Others have previously used ABMs to explore the mechanisms of disease progression in many conditions, including lung infections^87–91^ and cancer^92–95^. Our group has previously used ABM to study mechanisms of angiogenesis and vascular remodeling^46, 49, 96, 97^. For example, Walpole et al. developed an ABM of retinal angiogenesis in which endothelial cells migrate along a network of VEGF-secreting astrocytes to form new sprouts in the presence or absence of pericytes. Incorporation of pericytes in the model significantly improved the accuracy of the model’s prediction of microvascular remodeling, when experimentally validated against retinas from post-natal day 3 mice. While that model did not explicitly simulate intracellular signaling, it highlighted the importance of including both endothelial cells and pericytes in computational models of microvascular remodeling, as their interactions are key regulators of this process. Our multiscale model, which incorporates intracellular signaling in both cell types, extends the model by Walpole et al., by suggesting differential phenotypic responses by simulated ECs and pericytes to the same biomechanical and biochemical cues. This further underscores the importance of including both pericytes and ECs in models of microvascular remodeling and simulating not only their interactions with one another, but also their unique responses to the microenvironment which are interpreted by intracellular signaling networks that are unique to each cell type. Given that logic-based ordinary differential equations modeling has been used to represent many different cell types, including endothelial cells^51^, smooth muscle cells^98^, fibroblasts^52, 99–103^, and cardiomyocytes^53, 104–106^, our multiscale computational model can be extended to study other cell or tissue phenotypes by integrating new and previously established intracellular signaling models.

There are several limitations to our experimental studies and to our multiscale computational model. Overall, it is important to appreciate that IPF has a large amount of variability between patients and within different regions of the lung of the same patient. Therefore, the effects of nintedanib are likely to be equally variable, and while our findings align with prior studies, we capture only a few possible scenarios, and more research is warranted to fully appreciate nintedanib’s effects on the microvasculature. A limitation of our experimental explant model is the use of healthy mouse lung tissue, which may not exhibit the same phenotypic responses as diseased human lung. Despite this limitation, the experimental system provided a reproducible way to determine how ECM stiffness affects microvessel remodeling. The computational model only included ECs and pericytes, while in reality there are many cells in this interstitial niche, including epithelial cells, fibroblasts, and immune cells. Future versions of this model can incorporate epithelial cells to better capture the capillary-alveolar interface or fibroblasts, which contribute to the progressive stiffening of the matrix in IPF. Additionally, the model simulated only a small portion of lung (500µm x 500 µm) over a short time period of one week. The simulation size was largely limited by the number of cells, as more cells increased the overall time to compute. Additionally, the signaling pathways included in the EC and pericyte network models were limited to those that were deemed essential to represent EC-pericyte coupling and cell fate decisions. While it is likely that other signaling pathways are also important, and the current pathways in the models could be represented in more detail by including additional nodes, these adaptions can be included in future model iterations. Moreover, the independent validation we performed by comparing predictions to published literature suggested that the pathways that were included in the current version of the models were sufficient to invoke the major phenotypic states of ECs and pericytes. That being said, our review of the literature also revealed holes that can be filled by future studies.

Future studies should prioritize adapting the multiscale model to patient-specific contexts. For example, here we begin with an IF image of healthy lung vasculature to account for EC-pericyte ratios and a general vascular network starting point; but this can be replaced with other images in which specific EC and pericyte locations can be identified. Additionally, growth factors are known to be highly dysregulated in IPF, with evidence of extremely low VEGF availability^36^, increased abundance of PDGF^34^, and a decrease in Ang 1 associated with an increase in its agonist ligand, Ang 2^32^. Read-outs of these protein levels as serum biomarkers could allow for patient specific modeling. Moreover, incorporation of the other FDA approved drug for IPF, pirfenidone, a small molecule inhibitor of TGFβ, would also be of interest, as TGFβ regulates both pro-angiogenic and pro-quiescent pathways in ECs and pro-quiescent and pro-fibrotic pathways in pericytes^107, 108^. Additionally, here we assume the same stiffness is maintained for the entire time course; but future studies should model progressive stiffening as actually occurs in IPF. Lastly, other potential pathways for preserving EC-pericyte coupling can be explored to identify novel therapies and combination therapies to maintain microvessel homeostasis in IPF.

In conclusion, we have developed a multiscale modeling framework that integrates complex intracellular signaling networks in different cell types with cell-cell and cell-environment communication to elucidate mechanisms underpinning the loss of microvascular homeostasis in the context of IPF. This work highlights the need for more research on vascular remodeling in IPF and the benefits of combining computational and experimental models to study interacting heterogeneous cell populations in complex, remodeling microenvironments.

## Methods

### Logic Based Network Models

Logic-based network models are a form of ordinary differential equation-based mathematical models that have been used by us and others to represent intracellular signaling dynamics^45, 50, 109, 110^. For our multiscale model, we developed two independent logic-based network models, one for endothelial cells and one for pericytes. Both models were constructed based on data from an in-depth literature review of pathways that regulate EC-pericyte coupling and cell fate decisions, such as proliferation or migration. The MATLAB based software “Netflux” was used to develop both logic-based ordinary differential equation (ODE) based network models ^50, 109^. In order to achieve this, a normalized Hill ODE was implemented to represent signaling activity, utilizing logic gating to incorporate crosstalk. The default parameters for each species (mRNA, protein, cell process) are set to include a time constant ꞇ, an initial activity of 0 (y_int_), and a maximum activation of the species at 1 (y_max_). To account for the reaction time for different species the time constant parameter ꞇ is scaled for each species with signaling reactions requiring 6 minutes, transcription reactions requiring 1 hour, and translation reactions requiring 10 hours. Three additional parameters are used in each reaction in the model: reaction weight (w = 1), the Hill coefficient (n = 1.4), and EC50 (EC50 = 0.6). The excel sheet containing the constructed models can be found in the Supplementary Materials. The model outputs were visualized using Cytoscape^111^.

A sensitivity analysis was performed by knocking down each node, one at a time, and predicting the change in activity for all nodes in the network when compared to the baseline node activation. The baseline node activation, considered the control, was performed by simulating the model at the default values for each species and reaction parameter. Then one at a time, each node was knocked down (y_max_ = 0) and the activity for all nodes at steady state was recorded. The baseline activation levels for each node were then subtracted to calculate the change in activity.

### Endothelial Cell Logic Based Model

The endothelial cell logic-based network model consists of 34 species, 32 reactions, and 3 possible cell phenotypic states (**Figure 1A**). The seven input nodes to the model capture communication with other endothelial cells through DLL4 and Notch signaling to determine tip versus stalk fate, communication with neighboring pericytes through Ang1-Tie2 and N-cadherin signaling, and environmental cues such as VEGF, FGF, TGFβ, and ECM stiffness. ECs sense stiffness through many different proteins which we combine as one general integrin. The model predicts that the simulated cell will reside in one of three cell phenotypic states that represent the three high-level processes of vascular homeostasis and remodeling: angiogenic apoptotic, and quiescent.

In order to validate the EC logic-based network model, we used the model to simulate six different scenarios key scenarios for the multiscale model: 1) high VEGF, 2) low VEGF, 3) presence of a pericyte, 4) high TGFβ, 5) DLL4 stimulation from a neighboring EC, and 6) high FGF and the model predicted the EC phenotypic states (angiogenic, quiescent, and apoptotic) for each scenario. Published data from peer-reviewed papers that experimentally tested each of these scenarios were compared to the computational model predictions. A node was denoted as “increased” if the predicted model output was more than 5% higher than the baseline model output. A node was denoted as “decreased” if the predicted model output was more than 5% lower than the baseline model output. If the output after the perturbation differed from the baseline output by less than 5%, it was characterized as “no change”. The network model predictions matched the literature-reported experimental outputs for twelve out of the fourteen different scenarios that were simulated, or for 85% of the scenarios (**Figure 1C**).

### Pericyte Logic Based Model

The pericyte logic-based network model consists of 32 species, 34 reactions, and 3 possible cell phenotypic state outcomes (**Figure 1B**). There are five inputs to the pericyte model, two of which represent signaling with neighboring endothelial cells through Jagged 1 and Notch-mediated signaling, as well as through PDGF-BB-mediated paracrine signaling for pericyte recruitment. The other three inputs represent environmental cues, FGF, TGFβ and ECM stiffness, that have been shown to regulate pericyte fates in fibrotic environments. The three cell phenotypic states represent both homeostatic pericyte behavior, as defined by quiescence and migration, as well as the potential for myofibroblast transition through the expression of αSMA and collagens, as has been previously reported in the context of IPF^38, 39^.

Validation of the pericyte logic-based network model was performed using the same method as described above for the EC logic-based network model. Outputs were observed for the three phenotypic fate decisions for pericytes (migration, quiescence, and pericyte-to-myofibroblast transition (PMT)), in eight different scenarios: 1) high PDGF, 2) presence of an EC, 3) high ECM stiffness, 4) high FGF & EC presence, 5) high TGFβ, 6) TGFβ inhibition, 7) TGFβ inhibition & high ECM stiffness, and 8) PDGF inhibitor. Computational predictions for these scenarios were compared to experiments reported in published papers that evaluated the same scenarios. The same thresholds for “increased”, “decreased”, and “no change” that were used for validating the EC logic-based network model for were used for validating the pericyte model. The network model prediction matched the literature-reported experimental outputs for nine out of nine (100%) of the scenarios (**Figure 1D**).

### Evaluating Sub-Networks that Regulate Cellular Response to Nintedanib Treatment

To identify the nodes that regulate the effect of nintedanib on ECs and pericytes we performed a sensitivity analysis in the presence and absence of nintedanib. Knocking down each node in the absence of nintedanib describes the role of that node in nintedanib treatment. Comparing the difference in node activity in the untreated control compared to the activity in the presence of nintedanib describes the role of that node in regulating nintedanib’s effect on EC and pericyte cell fate. The nodes that have a differential effect dependent on nintedanib are identified as the sub-network that can be used as a mechanistic description of nintedanib activity. The sub-network analysis is performed in MATLAB then visualized used Cytoscape^111^.

### Agent-Based Model (ABM)

A two-dimensional (2D) ABM was built in NetLogo^54^, a freely available ABM software, representing an 500 μm x 500 μm slice of lung tissue. The approximate number of and positioning of ECs and pericytes was imported from an immunofluorescent image. The 2D ABM simulation space was discretized into an (x,y) grid of square pixels, each representing 12 μm x 12 μm. Each time step in the model represents 6 hours, and the total simulation time is one week, which is similar to timesteps that have been used in ABMs published by our lab and has proven to be computationally tractable^40, 47, 112^.

Behaviors of simulated pericytes and EC “agents” are dictated by the outputs from the logic-based network models for each cell type. At each time step, every agent records its environmental cues (i.e., stiffness, proximity to surrounding agents, cytokines, etc.) and uses them as the inputs to the logic-based network models, which in turn, predict cell behaviors (i.e., quiescence, migration, apoptosis, etc.). The agent then enacts that behavior, and the process repeats on the next timestep.

Based on published literature that has characterized the relative stiffnesses of different regions of tissue in IPF lungs^42^, three different environmental stiffness levels representing healthy (2 kPa), immature fibrosis (10 kPa), and mature fibrosis (20 kPa) were simulated.

### Connecting the Models Across Scales

The general principles for connecting the logic-based models to the ABM are based on a previously published multiscale model from our lab^45^. In the ABM cellular agents can interact with each other and the extracellular environment and this information will be stored in the ABM according to the type of interaction. These variables will then be translated to a normalized scale for input to the logic-based network model. The network model is then run until steady state is achieved and the steady state values for each cellular phenotype are input to the ABM for each cell (**Figure 9**). The ABM is scaled such that each time step, or tick, is 6 hours and is run for a total of 28 ticks, or 7 days, to capture short term microvascular remodeling. At each time step each agent will record growth factor levels in the environment, presence of neighboring cells, and ECM stiffness then translate them into relative activation levels for each input of the network model. The cell will call its respective network model based on if it is an EC or a pericyte, the model will be executed in Python, then return values for cell fates in the form of relative activations to the ABM. These will be rounded to the nearest hundredth and the cell agent in the ABM will take on the cell fate of the node that is most highly activated.

**Figure 9.**
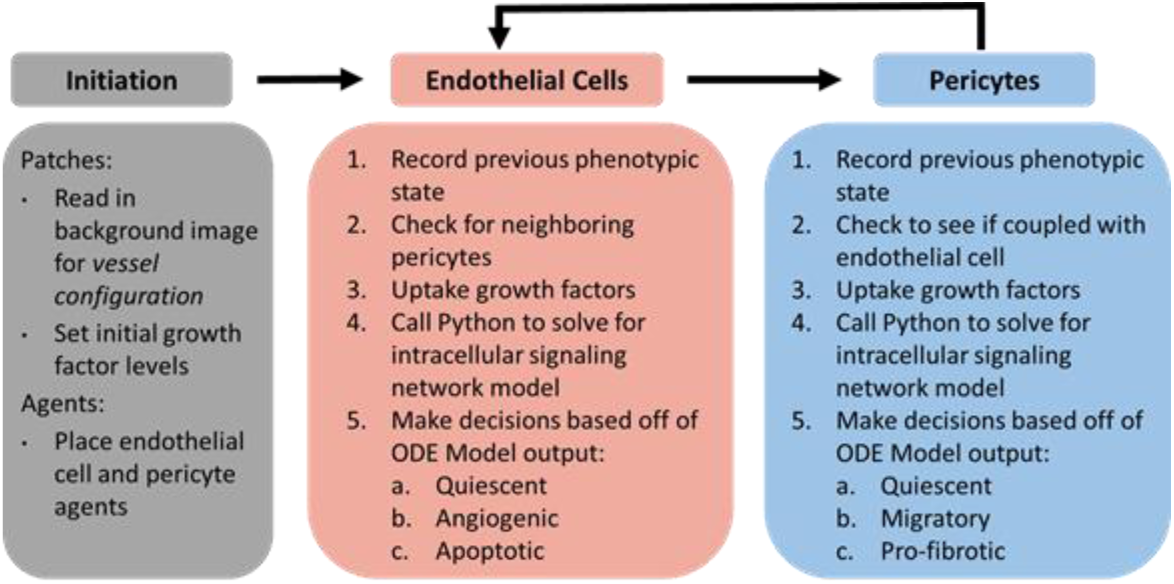
Flow chart of multiscale model work flow. At the initiation of a model the background configuration is set up from an IF image and the initial level of each growth factor is set according to user input. Agents are then placed in specific locations based on IF image data. Then at every time step EC agents will first intake their surrounding environmental cues, input them into Python, and receive a cell fate decision that will then be executed in the ABM. Finally, pericytes repeat the same process as ECs: update cues, solve the ODE in python, then execute the returned cell fate decision. This cycle repeats until the model is terminated.

Occasionally, a tie will occur for nodes that are regulated by similar inputs. For ECs, in the case of a tie between the quiescent and angiogenic cell fates the agent will have a 1 in 100 chance of undergoing angiogenesis instead of remaining quiescent. For pericytes, if there is a tie between the migratory and PMT cell fates, there is a 1 in 100 chance that the pericyte will undergo PMT.

Cellular interactions are stored as a binary, the cell type of interest either is or is not a neighbor of the primary agent. For ECs, there are two cellular interactions that are dependent on neighboring cells: 1) the presence of a neighboring pericyte and 2) the presence of a neighboring tip cell. Pericytes maintain both physical and paracrine connections to ECs and this is represented in the model through n-cadherin signaling and angiopoietin 1 (Ang-1) secretion. If the EC is in the presence of a pericyte the input for pericyte n-cadherin variable and Ang-1 variable will have an input of 1. If a pericyte is not in the neighboring eight patches pericyte n-cadherin will be set to 0 and Ang-1 to 0.1 to account for potential secretion from other cells in the environment, although pericytes are the major source of Ang-1^113–115^. If the EC is next to another EC with the “angiogenic” state indicating that it will become a tip cell and is expressing DLL4, the variable for DLL4 will be set to 1 to encourage stalk cell fate, or a “quiescent” cell state^116, 117^. For pericytes their only cellular interaction is with ECs through jagged 1 – notch 3 signaling. If they are within eight patches of an EC, the input for EC jagged 1 will be set to 1, otherwise this will be set to 0^15, 118^.

Extracellular interactions include both integrin mediated response to ECM stiffness and signaling through growth factor receptors. Cellular response to ECM stiffness is scaled according to the observed stiffnesses of fibrotic lung^42^ with the maximum relative activation of the node representing 20 kPa. As previously described in Rikard et al.^45^, The relative activation of each growth factor receptor is translated from literal pg/mL values in the ABM using a Hill equation where the dissociation constant of the receptor, K_d_, is equivalent to the ligand concentration at which 50% of the receptors are activated. There are four growth factors in this model that are represented using Hill equations: 1) TGFβ (equation 1), 2) FGF-2 (equation 2), 3) VEGF-A (equation 3), and 4) PDGF-BB (equation 4). The K_d_ and concentration at which 50% of the receptors are activated by ligand are listed in **Table 1**.

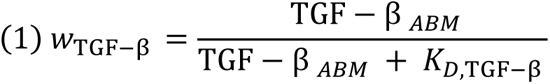

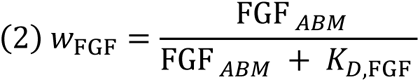

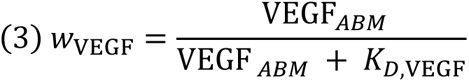

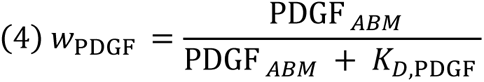

**Table 1.** Dissociation constant values used in multi-scale computational model.

| Growth Factor | K<sub>d</sub> (mM) | Concentration for 50% Activation(pg/mL) | Source |
| --- | --- | --- | --- |
| TGF-β | 0.028 | 700 | 45, 124 |
| FGF-2 | 62 | 1116000 | 125, 126 |
| VEGF-A | 0.15 | 4056.3 | 127 |
| PDGF-BB | 0.44 | 10692 | 128 |

All initial inputs to the network model are set to 0.35 unless they are dependent on a conditional relationship, such as ECM, or presence of a neighboring cell. These conditional values are listed in **Table 2**. To capture heterogeneity in growth factor availability the microenvironment each patch in the model multiplies its initial levels of each growth factor by a random factor ranging from 0.5 to 1.5. Additionally, when a cell uptakes a growth factor, it will remove up to 5% of the value it absorbed from the patch to account for the continuous background secretion of growth factors into the microenvironment. The value for ECM stiffness is not changed by any cellular behaviors.

**Table 2.**
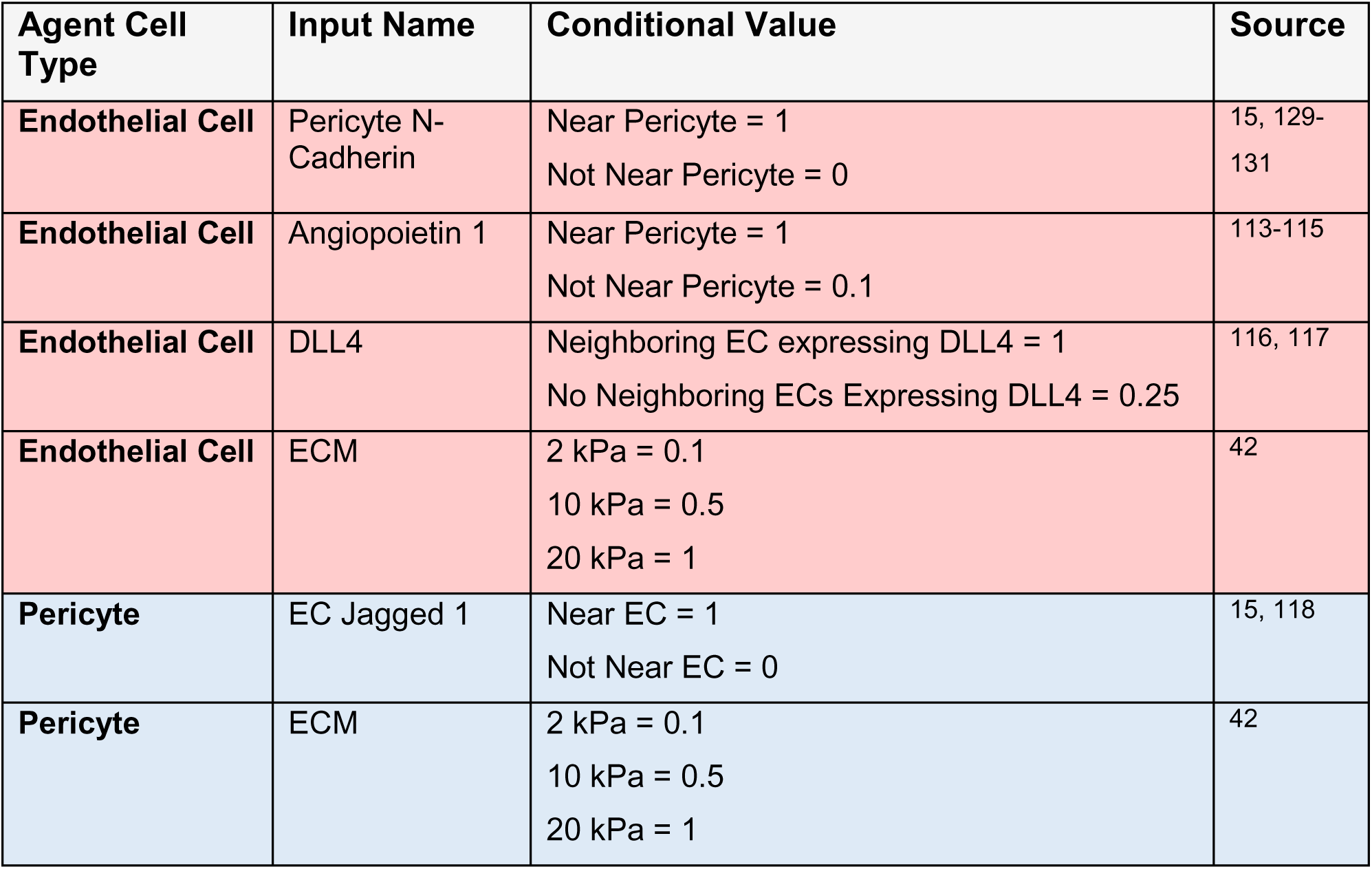
Conditions for different input values according to cellular or extracellular interactions in the ABM.

Nintedanib and YAP/TAZ inhibition was modeled as by inhibiting the weight, *w*, of the reaction producing the node of interest. For example, the input weight for the equation representing VEGF to VEGFR binding was reduced from 1 to 0.1. All ABM simulations are conducted in NetLogo^54^ which then accesses Python via the “Py” extension to run the logic-based network models. This allows each agent to independently enter its inputs into the logic-based network model.

### Mice

All procedures described in this study were performed in in accordance with the Institutional Animal Care and Use Committee of the University of Virginia. *Myh11*-CreER^T2^ ROSA floxed STOP tdTomato mice were bred from *Myh11*-CreEr^T2^ mice (cat. no. 019079; The Jackson Laboratory, Bar Harbor, ME) and ROSA floxed STOP tdTomato mice (007914; The Jackson Laboratory), on C57BL/6J background. Expression of tdTomato in cells of the myosin heavy chain 11 (Myh11) lineage (including pericytes and vascular smooth muscle cells) was induced by feeding 6 to 8 week old male mice a tamoxifen-containing diet (Envigo) for 14 days^39, 72, 119, 120^. Only male mice were used in this study, as the Cre gene cassette was inserted on the Y chromosome. Experiments were initiated 2 weeks after the mice were returned to a normal chow diet to allow for clearance of tamoxifen and to reduce the possibility that cells that transiently expressing Myh11 would be falsely identified as a pericyte.

### PEGylation of peptides and growth factors

PEG hydrogels were functionalized with the following peptides and proteins purchased from Genscript (Piscataway, NJ): RGDS, a fibronectin-derived adhesion ligand, GGGPQGIWGQGK (PQ), a protease-sensitive degradable linker, basic fibroblast growth factor (FGF-2), an angiogenic endothelial cell mitogen, and platelet-derived growth factor (PDGF) which also contributes to angiogenic signaling in ECs and pericytes. RGDS and PQ were conjugated to 3.4 kDa acrylate-PEG-succinimidyl valerate (PEG-SVA, Laysan Bio, Arab, AL) according to methods described previously^121^. PEG-SVA and peptides were dissolved in dimethyl sulfoxide (DMSO, Sigma) at a 1.1:1 (RGDS:PEG-SVA) and 2.1:1 (PQ:PEG-SVA) molar excess, with a 2:1 molar ratio of N,N-diisopropylethylamine (DIPEA, Sigma-Adrich, St. Louis, MO) to PEG-SVA for 24 hours with constant agitation to form acrylate-PEG-RGDS (PEG-RGDS) and acrylate-PEG-PQ-PEG-acrylate (PEG-PQ). Conjugates were dialyzed against ultrapure water overnight, lyophilized, and their purity and conjugation efficiency were assessed via gel permeation chromatography (GPC, EcoSEC Elite, Tosoh Biosciences, King of Prussia, PA) with refractive index (RI), ultraviolet (UV), and multi-angle light scattering (MALS) detectors. PEG-PDGF and PEG-FGF were formed by conjugation of PEG-SVA to PDGF and FGF at a 1:200 molar ratio in sterile 50 mM sodium bicarbonate (pH 8.5) buffer at 4 °C for 4 days followed by 4 hours at ambient temperature. Growth factor conjugation was verified through SDS-PAGE.

### Fabrication of hydrogels of varying stiffness

The stiffness of 5% (w/v) PEG-PQ hydrogels, which are characterized as having an elastic modulus of approximately 20 kPa^122^, can be tuned independently of polymer density through the incorporation of a vinyl-bearing crosslinking inhibitor as described by Chapla et al^123^. This method was used to fabricate gels having softer moduli through the addition of an alloc-protected lysine at a ratio of 2:1 alloc:acrylate (to form 10 kPa hydrogels) and 3.5 alloc:acrylate (to form 2 kPa hydrogels) to hydrogel precursor solutions. Precursor solutions were prepared in 1 mL sterile HEPES-buffered saline (HBS, pH 7.4) from 50 mg/mL PEG-PQ, 3.5 mM PEG-RGDS and the alloc-protected lysine in HBS, then sterile-filtered using a 0.2 µm syringe filter (Whatman Puradisc Regenerated Cellulose Syringe Filter, Cytiva). To each precursor solution, 0.7 nmol/mL PEG-PDGF and 0.27 PEG-FGF were added as matrix-bound angiogenic factors. Hydrogels were UV-cured in silicone molds (height = 1 mm, ID = 6 mm) adhered to the bottom of 24-well glass bottom plates functionalized via pre-treatment with ethanol containing 2% (v/v) 3-(trimethoxysilyl)propyl methacrylate for 4 days to methacrylate the glass in each well. 35 μl of PEG precursor solution containing PEG-RGDS, PEG-PDGF, PEG-FGF, and 10 μL/mL of the photoinitiator 2-dimethoxy-2-phenylacetophenone (acetophenone; Sigma-Aldrich), prepared by dissolving 300 mg acetophenone in 1 ml of N-vinylpyrrolidone (NVP), was pipetted into the center of each mold, and the plate exposed to UV light (365 nm, 10 mW/cm^2^) for 30 s. The gels were allowed to swell in PBS overnight and then incubated in culture media.

### Lung Explant Assay in Engineered Hydrogel

We performed a lung explant assay as previously described^72^. After the tamoxifen clearance period, when mice were approximately 12-16 weeks of age, three mice were humanely euthanized via carbon dioxide asphyxiation. The chest of the mouse was opened such that the heart and lungs were visible and a cardiac perfusion was performed using 10mL of phosphate buffered saline (PBS). The heart and lungs were then removed together, and the lungs were surgically dissected from the heart and surrounding connective tissue. Lungs were placed in cold PBS on ice until they were sectioned into strips of lung tissue for the explant assay. Each lobe was placed on the tissue slicer (Stoelting) and cut into 1mm long strips. These strips were then turned 90° and sliced again to yield 1 mm x 1 mm explants. Each explant was then placed into a well of a 96-well plate containing cold PBS until plating in the engineered hydrogel.

On Day 0, one explant strip was placed in the center of each hydrogel contained within a tissue-culture well of a 24 well plate. Each well received 500µL of Endothelial Growth Media 2 – Microvascular Formula (EGM2-MV), 250µL of which was replaced every other day. In total, six 24-well plates each contained hydrogels with three different stiffnesses (2 kPa, 10 kPa, and 20 kPa) with 6 wells per stiffness level.

### Explant Assay Imaging

Explants were imaged on Day 7, and Day 14. On Day 7, five plates were imaged using brightfield imaging on the Leica THUNDER Imager using the 10X objective (Leica). Images were then sorted into four groups, as previously described^72^, based on the identification of organized vessel-like structures versus disorganized cell infiltration. The number of images in each category were averaged in each well and then across wells of the same condition in a plate. This accounts for variability between explants based on the region of the lung from which they were isolated. Each plate was then considered an independent sample, *N*. The percent of images containing vessel sprouts for each experimental condition (2kPa, 10kPa, and 20kPa) was calculated.

On Day 14 all six plates were imaged again using the Leica THUNDER Imager with brightfield on the 10X objective. These were then analyzed based on the amount of organized cellular infiltration into the surrounding gel from the tissue explant. In a blinded analysis, this infiltration was separated into five categories from no evidence of vessel infiltration (Category 1) to extensive vessel-like cellular infiltration that fills the image frame (Category 5). **Table 3** shows representative images of these different categories. Groups were analyzed using the same method as the Day 7 calculations with each independent plate representing a sample, *N*.

**Table 3.**
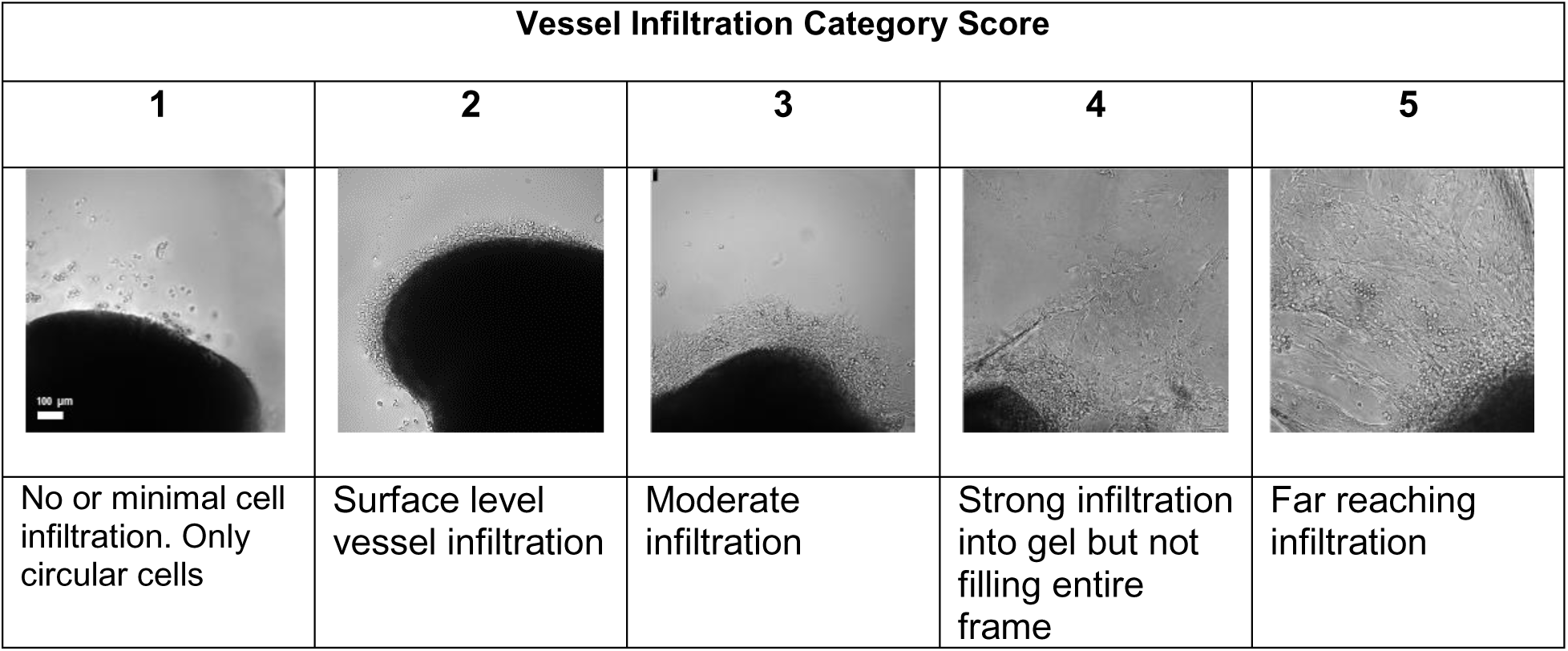
Guidelines for vessel infiltration category score.

| Vessel Infiltration Category Score |  |  |  |  |
| --- | --- | --- | --- | --- |
| 1 | 2 | 3 | 4 | 5 |
| No or minimal cell infiltration. Only circular cells | Surface level vessel infiltration | Moderate infiltration | Strong infiltration into gel but not filling entire frame | Far reaching infiltration |

### Treatment of Lung Explants with Nintedanib

After Day 7 imaging was completed the six plates were split into two groups: three plates received nintedanib and three plates received DMSO as a control. The nintedanib was prepared fresh for each media change at a concentration of 50nM in 0.05% DMSO in EGM-2MV. The control groups received 0.05% DMSO in EGM-2MV.

### Statistical Analyses

For the computational models a One-way ANOVA was used to compare between groups in different simulated stiffnesses with a Tukey’s post-hoc test. For the nintedanib treated simulations the nintedanib treated were compared to their respective control group using an unpaired one-tailed student’s t-test. For the experimental results statistics was performed as previously described^72^. In short, each 24-well plate was considered an independent sample, N, and all images acquired for that plate were averaged for each experimental sub-group. For comparing groups of different stiffness in experimental test 1, a One-way ANOVA with Tukey’s post-hoc test was used. For experimental test 2, each nintedanib treated sample was compared to its DMSO treated control using a Student’s t-tests were used for statistical analyses between groups with the same stiffness. Statistical significance is asserted at p-values < 0.05. All data are presented as average +/- standard deviation.

## Supporting information

EC Netflux Model

Pericyte Netflux Model

## Acknowledgements

The authors would like to thank Riley Hannan, Ph.D. for his assistance with collecting the IF image used in the ABM setup.

## Author Contributions

Figure and manuscript preparation: JLD, SMJA, DJC, LJT, and SMP. Conceptualization: JLD, THB, CAB, JJS, LJT, and SMP. Study design: JLD and SMP. Data collection: JLD, SMJA, DJC, TNT, and ACB. Data analysis: JLD, SMJA, DJC, and TGE. All authors participated in the review and approval of the final manuscript.

