## Supplementary Figures for "Multiscale computational model predicts how environmental changes and drug treatments affect microvascular remodeling in fibrotic disease"

*Supplementary Materials*

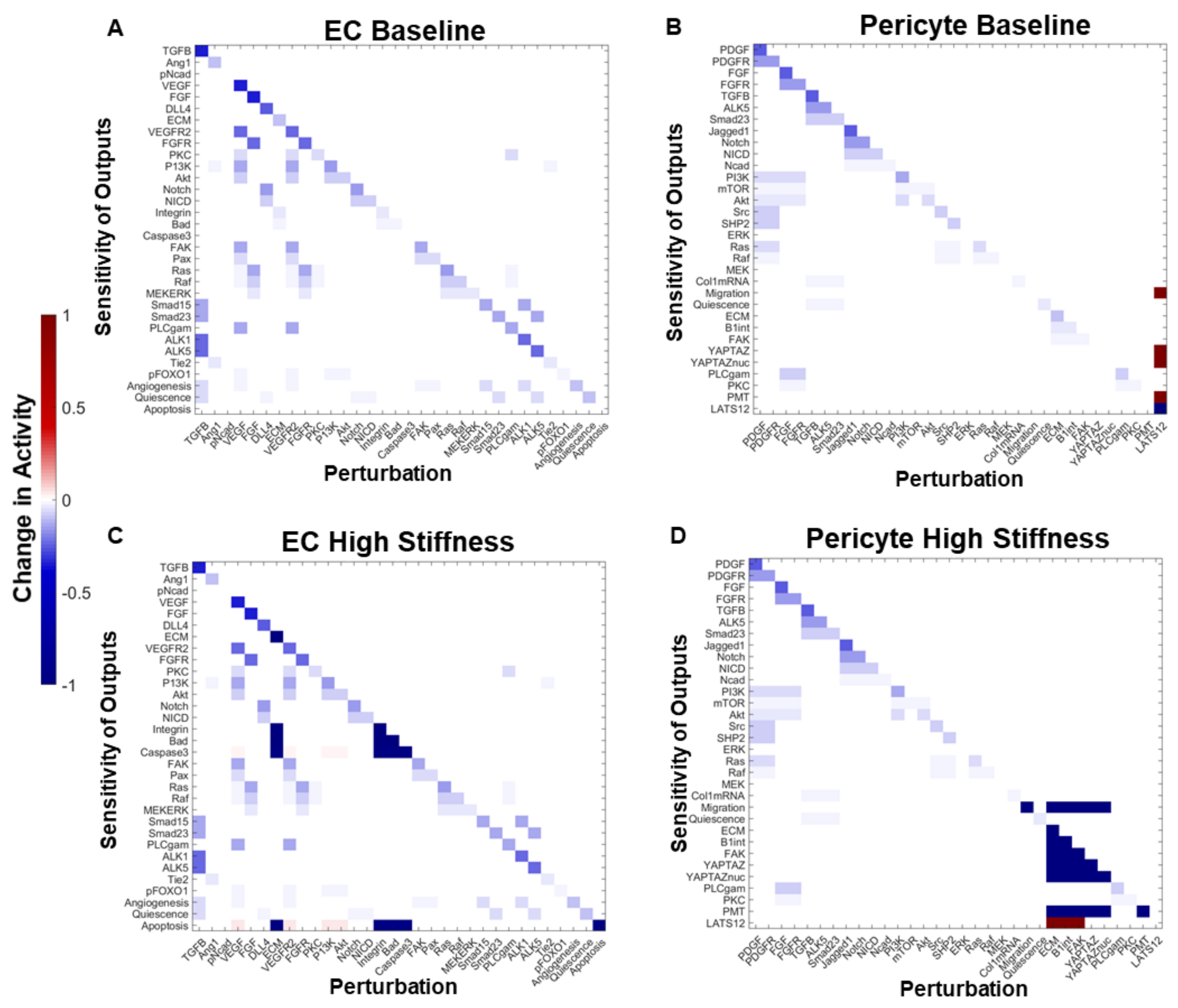


**Figure S1. Sensitivity analysis of EC and pericyte network models at baseline and in response to increased mechanical stiffness.** (A) Sensitivity analysis of EC network model confirms the model is not overly sensitive to any one set of nodes. (B) Sensitivity analysis on pericyte network model identifies sensitivity in the model to LATS1/2 KO. (C) Sensitivity analysis in the presence of high mechanical stiffness $w_{stiffness}$ = 1 demonstrates sensitivity in integrin mediated stiffness sensing. (D) Sensitivity analysis in the presence of high mechanical stiffness $w_{stiffness}$ = 1 demonstrates sensitivity in to integrin mediated stiffness sensing.

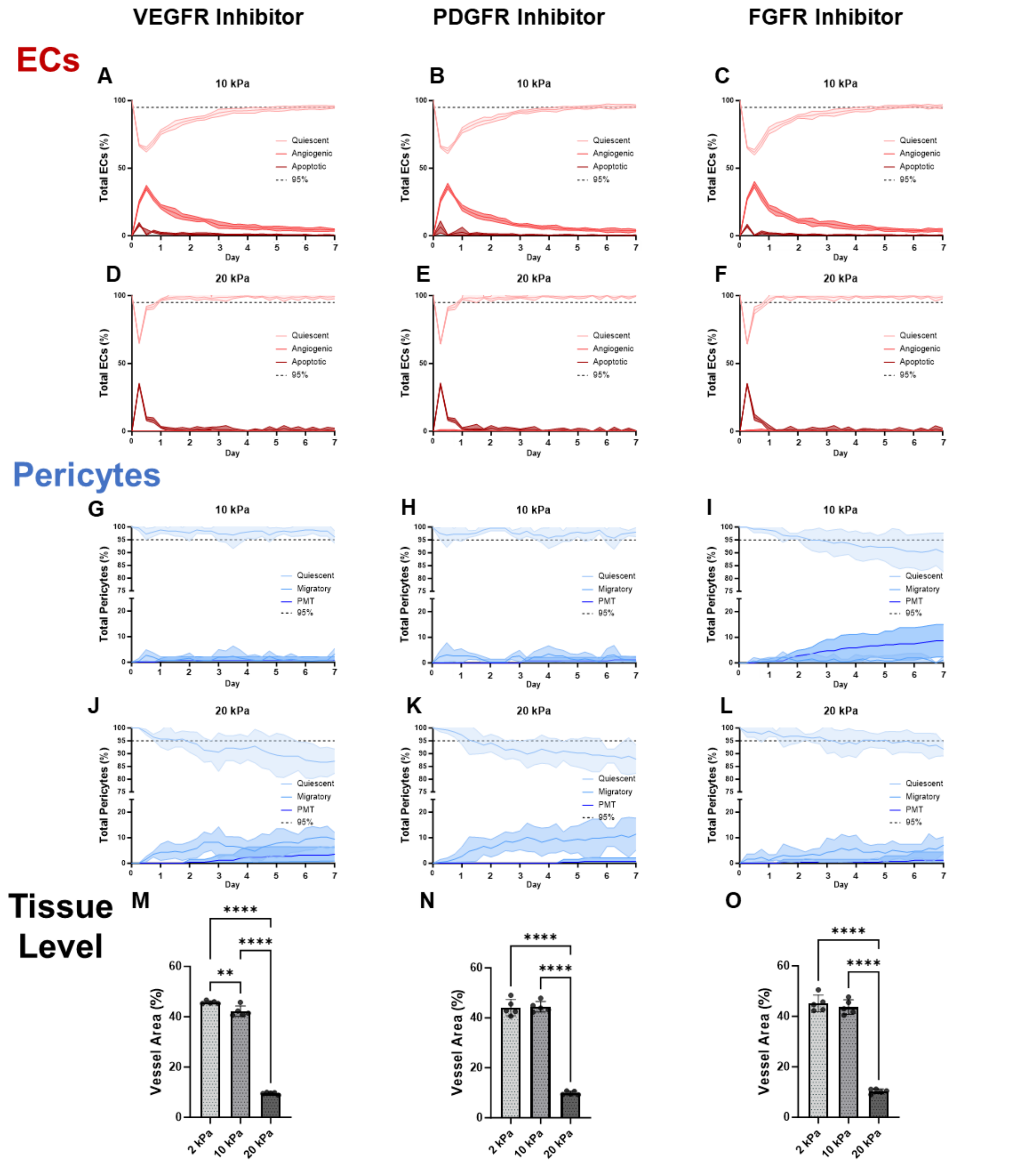


**Figure S2. EC response to single receptor inhibition in fibrotic foci (A-F).** (A) At 10 kPa treatment with a VEGFR inhibitor delays 95% EC quiescence steady state recovery to Day 6. (B) Treatment with a PDGFR inhibitor causes a recovery of the 95% quiescent EC steady state at Day 5. (C) Treatment with FGFR results in a recovery of 95% EC quiescent steady state at Day 5 with lower levels of apoptosis than PDGFR inhibition. (D) At 20 kPa, a sharp decline in 95% quiescent EC steady state is recovered by Day 1 and associated with a spike in apoptosis with treatment with a VEGFR inhibitor. (E) When treated with a PDGFR inhibitor, a sharp decline in 95% quiescent EC steady state at 20 kPa is recovered by Day 1 and associated with a spike in apoptosis. (F) At 20 kPa, a sharp decline in 95% quiescent EC steady state is recovered by Day 1 and associated with a spike in apoptosis when treated with an FGFR inhibitor. **Pericyte response to single receptor inhibition in fibrotic foci (G-L).** (G) Pericyte 95% quiescence levels are maintained in the presence of a VEGFR inhibitor alone. (H)

PDGFR inhibition maintains the average pericyte quiescence level above the 95% steady state threshold. (I) Treatment with an FGFR inhibitor leads to a sharp decline of pericyte quiescence an increase in PMT at Day 1 that does not reverse. (J) VEGFR inhibition leads to a sharp decline in pericyte quiescence at Day 1 that is not recovered followed by increased levels of migration and PMT starting at Day 2. (K) PDGFR inhibition leads at 20 kPa leads to a sharp decline in pericyte quiescence at Day 1 that is mainly associated with an increase in migration. Evidence of PMT is not observed until Day 4. (L) At 20 kPa treatment with an FGFR inhibitor delays the loss of pericyte 95% quiescent steady state until Day 3 and it is briefly recovered on Day 5 before declining with an associated rise in PMT. **Tissue level response to single receptor inhibition (M-O)** (M) Significant decrease in vessel area in response to increasing stiffness even with VEGFR inhibitor treatment. (N) Treatment with a PDGFR inhibitor prevents a significant decrease in vessel area at 10 kPa but not 20 kPa. (O) Treatment with a PDGFR inhibitor prevents a significant decrease in stiffness at 10 kPa but not 20 kPa. *Statistics: One-Way ANOVA with Tukey's post-hoc test for multiple comparisons, N = 5, error bars represent standard deviation,  shaded regions represent 95% confidence interval*

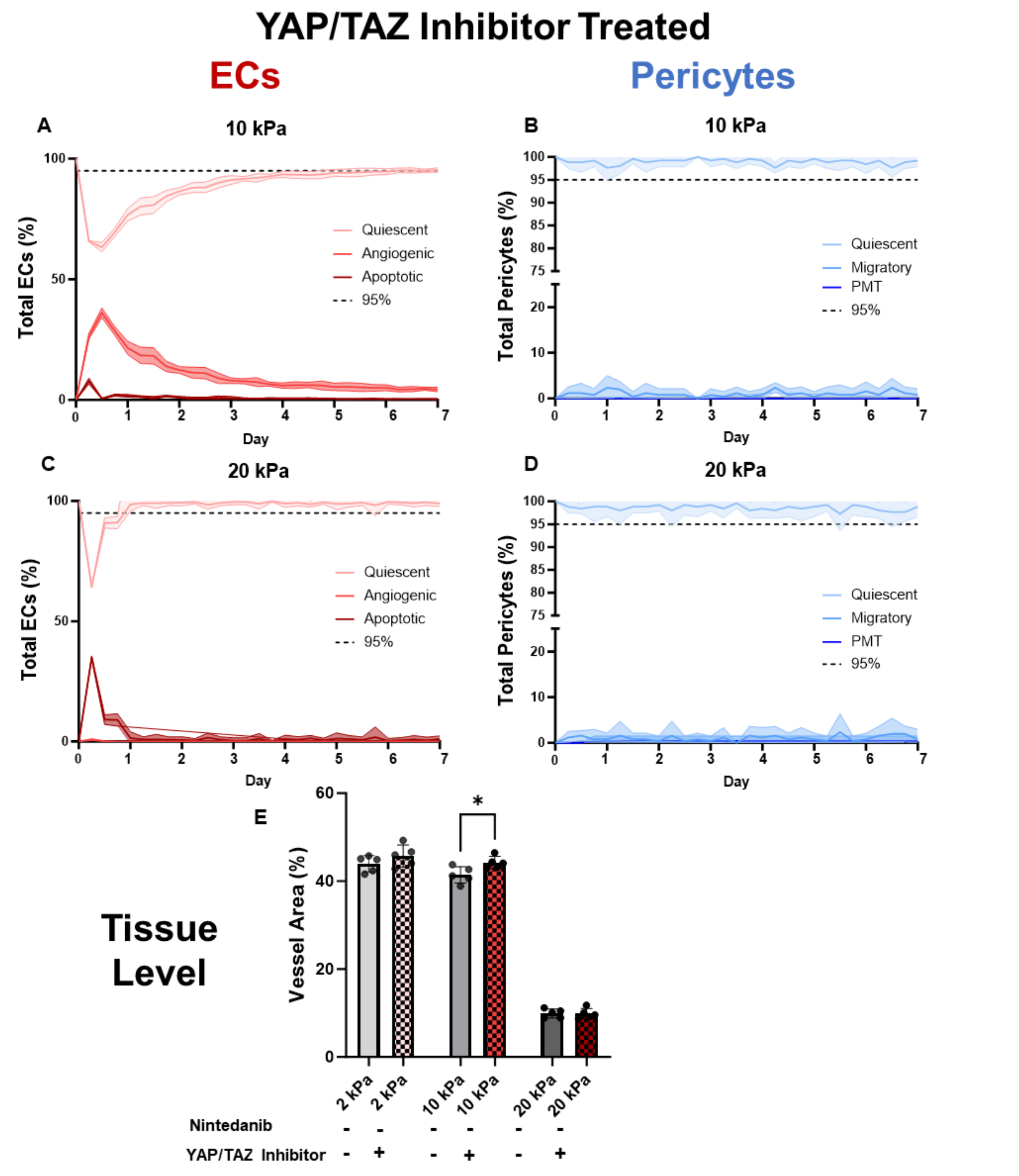


**Figure S3. The effect of YAP/TAZ inhibition alone on EC-pericyte coupling and vessel area** (A) A decline in the percent quiescent ECs is associated with increases in the percent of ECs that are angiogenic or apoptotic and eventually recover 95% quiescent cells at Day.6 (B) YAP/TAZ inhibition maintains over 95% of pericytes in the quiescent state at 10 kPa throughout the time course. (C) An initial decrease in the percent of quiescent ECs is accompanied by a large spike in apoptotic ECs. Although 95% quiescence is recovered on Day 1. (D) At 20 kPa, YAP/TAZ treatment alone preserves pericytes quiescence levels above 95%. (E) Vessel area is significantly increased when compared to nintedanib treated alone at 10 kPa, but not at 2 or 20 kPa. *Statistics: Two-tailed t-test, N = 5, error bars represent standard deviation, shaded regions represent 95% confidence interval*
